# Investigating the oral microbiome in retrospective and prospective cases of prostate, colon, and breast cancer

**DOI:** 10.1101/2022.10.11.511800

**Authors:** Jacob T. Nearing, Vanessa DeClercq, Morgan G.I. Langille

## Abstract

The human microbiome has been proposed as a useful biomarker for several different human diseases including various cancers. To answer this question, we examined salivary samples from two Canadian population cohorts, the Atlantic Partnership for Tomorrow’s Health project (PATH) and Alberta’s Tomorrow Project (ATP). Sample selection was then divided into both a retrospective and prospective case control design examining individuals with prostate, breast, or colon cancer. In total 89 retrospective and 260 prospective cancer cases were matched to non-cancer controls and saliva samples were sequenced using 16S rRNA gene sequencing to compare bacterial diversity, and taxonomic composition. We found no significant differences in alpha or beta diversity across any of the three cancer types and two study designs. Although retrospective colon cancer samples did show evidence on visual clustering in weighted beta diversity metrics. Differential abundance analysis of individual taxon showed several taxa that were associated with previous cancer diagnosis in all three groupings within the retrospective study design. However, only one genus (*Ruminococcaceae UCG-014*) in breast cancer and one ASV (*Fusobacterium periodonticum*) in colon cancer was identified by more than one differential abundance (DA) tool. In prospective cases of disease three ASVs were associated with colon cancer, one ASV with breast cancer, and one ASV with prostate cancer. None overlapped between the two different study cohorts. Attempting to identify microbial signals using Random Forest classification showed relatively low levels of signal in both prospective and retrospective cases of breast and prostate cancer (AUC range: 0.394-0.665). Contrastingly, colon cancer did show signal in our retrospective analysis (AUC: 0.745) and in one of two prospective cohorts (AUC: 0.717). Overall, our results indicate that it is unlikely that reliable oral microbial biomarkers of disease exist in the context of both breast and prostate cancer. However, they do suggest that further research into the relationship between the oral microbiome and colon cancer could be fruitful. Particularly in the context of early disease progression and risk of cancer development.

## Introduction

The oral microbiome is a highly diverse microbial community that is shaped by several different dietary, anthropometric and lifestyle choices (Belstrøm et al., 2014; Nearing et al., 2020). Recent works have shown that this community of microbes plays important roles in both oral and systemic health (Wade, 2013). For example, the composition of the oral microbiome has been associated with oral diseases such as periodontitis (Lundmark et al., 2019) or more distal diseases such as colorectal cancer (CRC) (Flemer et al., 2017). Recent work has proposed that the oral microbiome may possess biomarkers for the identification of various diseases. However, research on some of the most common cancers such as prostate, colon, and breast cancer are limited. In the year 2022, it is estimated that 28,900 cases of breast cancer, 24,600 cases of prostate and 24,300 cases of colon cancer will be diagnosed in the Canadian population (Brenner et al., 2022). While numerous works have previously examined the associations of these three cancers and the human microbiome most studies have focused on their relationship with the gut microbiota or microbiota associated with the organ of interest.

Of these cancers, prostate cancer has arguable received the least amount of attention within the microbiome field. Studies on the human microbiome and this disease have been mostly focused on prostate tissue, and the gastrointestinal tract (Javier-DesLoges et al., 2022). Investigation into the microbial inhabitants of prostate tissue have given mixed results due to the low biomass of these samples and contamination issues. Indeed, studies on healthy prostate tissue have given conflicting results, with some indicating the presence of bacteria and others finding no evidence of bacterial inhabitants (Porter et al., 2018). However, in prostate cancer multiple works have demonstrated evidence for bacterial communities in tumors and benign tissue (Cavarretta et al., 2017; Feng et al., 2019; Yow et al., 2017). This led to interest in determining whether specific microbes might be associated with tumor and non-tumor tissue. To date these studies have given inconsistent results about specific bacteria but ultimately point to similar overall microbial community structures between tumor and benign prostate tissue (Cavarretta et al., 2017; Feng et al., 2019; Yow et al., 2017).

In contrast to the prostate microbiome, it is well known that the gastrointestinal tract contains a rich population of microbes including bacteria, archaea, and fungi (Huttenhower et al., 2012; Wilson & Blitchington, 1996). Work on these communities have shown a myriad of associations with various diseases, metabolites, and immunological states (Petrosino, 2018). Several smaller studies have examined the relationship between the gut microbiome and prostate cancer with mixed results. Two studies by Liss et al., and Golombos et al., found higher levels of Bacteroidetes in patients with prostate cancer indicating potential linkages between the disease and this broad taxonomic group (Golombos et al., 2018; Liss et al., 2018). Furthermore, work by Matsushita et al., in Japanese men with high-Gleason prostate cancer linked an enrichment of short chain fatty acid producing bacteria in the gut and cancer status (Matsushita et al., 2021). However, other works by Alanee et al. and Katz et al., have found no significant differences between individual’s with and without prostate cancer (Alanee et al., 2019; Katz et al., 2022). Leading to mixed results on whether the microbial composition on the gut is truly linked to the development of prostate cancer.

Considering these investigations, work on the oral microbiome and prostate cancer is rather lacking. To the best of our knowledge only one study has examined this potential linkage despite previous works linking periodontitis with the likelihood of prostate cancer development (Lee et al., 2017; Wei et al., 2021). In 2017, work by Estemalik et al., found in a group of 24 patients with chronic prostatitis or benign prostatic hyperplasia that 70.8% had one or more bacteria found within their oral cavity to also be residing within prostatic secretions (Estemalik et al., 2017). These results may indicate a linkage between prostate inflammation and the oral cavity warranting further investigation into the linkage between the oral microbiome and prostate health.

Breast cancer is one of the most commonly diagnosed cancers in females and is one of three cancers examined within this report (Brenner et al., 2022). Studies on breast cancer and the human microbiome have shown variable results with some studies suggesting associations between disease status and microbial community composition within the gut, breast tissue, and urine (Fernández et al., 2018). Indeed, early work on the gut microbiota and breast cancer suggested the presence of an “estrobolome” (Järvenpää et al., 1980; Plottel & Blaser, 2011). Which considers the functional capacity of microbes to metabolize estrogen and estrogen related products within the gut. Evidence of the estrobolome was first uncovered during the 20^th^ century by several studies that identified fluctuations in estrogen and estrogen related metabolites within the plasma, urine, and feces of pregnant women using antibiotics (Järvenpää et al., 1980; F. Martin et al., 1975; Tikkanen, Pulkkinen, et al., 1973; Tikkanen, Adlercreutz, et al., 1973; Willman & Pulkkinen, 1971). Since these initial discoveries further work has characterized bacterial enzymes within the gut such as β-glucuronidase that can process conjugated estrogen metabolites (Ervin et al., 2019; Järvenpää et al., 1980). From these findings it has been suggested that the estrobolome may play a role in the risk of breast cancer development through the control of recirculating estrogen levels (Adams, 2016; Chen & Madak-Erdogan, 2016; Plottel & Blaser, 2011).

Investigation into this hypothesis through examining the relationship between the gut microbiome and breast cancer status has shown varying results. For example, work by Goedert et al, found reduced microbial diversity within the gut of post-menopausal breast cancer patients, however, work in 2018 by Zhu et al., reported significant findings in the opposite direction (Goedert et al., 2015; J. Zhu et al., 2018). Similarly, several gut microbes including those that possess β-glucuronidases have been associated with breast cancer status depending on study and menopausal status (Byrd et al., 2021; Goedert et al., 2015, 2018; Hou et al., 2021; J. Zhu et al., 2018). However, due to the high variability of results between studies and the lack of non-sequenced based validation, no strong conclusions have been made on the role of any specific gut taxon and breast cancer risk.

In addition to the gut, several studies have examined breast cancer status and microbial communities within and on breast tissue and urine. Although, these studies just like the gut, have also shown variable results indicating the need for further work within the field (Fernández et al., 2018). Interestingly, in the case of the oral microbiome, despite work showing that breast cancer is associated with periodontal disease to the best of our knowledge only two published studies have examined its relation with cancer status (Chung et al., 2016). Early work by Wang et al., examining oral rinses from individuals in the United States found no differences in community composition or specific taxa (H. Wang et al., 2017). However, recent work by Wu et al., did show differences in both microbial diversity and specific taxon within salivary samples from the Ghana Breast Health Study (Wu et al., 2022). These inconsistent results highlight the need for further investigation into the potential relationship of the oral microbiome and breast cancer in different populations.

Finally, one of the most well studied cancers in the context of the human microbiome is CRC. Numerous studies have been conducted on the relationship between CRC and the gut microbiome showing significant differences in community composition between cancer and non-cancer individuals (Zhao et al., 2021). Highlighted within these studies have been the relationships between various bacteria and CRC development or progression; including *Fusobacterium nucleatum*, enterotoxigenic *Bacteroides fragilis*, and (pks+) *Escherichia coli* (Garrett, 2019). However, to date it is still not entirely clear how influential these species are on the development or progression of the disease. For example, work by Dziubańska-Kusibab et al., has shown that colibactin produced by pks+ *E. coli* can cause DNA damage within the colon and is associated with mutational signatures often found in colon cancer patients (Dziubańska-Kusibab et al., 2020). However, (pks+) *E. coli* has yet to be formally recognized as a causative agent of CRC by the World Health Organization.

Barring from these investigations has been research on the relationship between the oral microbiome and CRC. Work in this area has shown that shifts in community composition within the oral microbiome has been associated with disease and in some cases may be predictable. Work in 2018 by Flemer et al., found that at diagnosis the oral microbiome could classify individuals with or without colon cancer with an area under the receiver operator curve (AUROC) of 0.91 (Flemer et al., 2017). Furthermore, their work along with others has also shown that many taxa commonly attributed to the oral cavity are enriched in the gut microbiome of CRC patients (Flemer et al., 2017; Thomas et al., 2019). Additionally, three further studies on the oral microbiome and colorectal cancer within various group settings reported distinct bacterial signatures associated with disease diagnosis (Komiya et al., 2019; Y. Wang et al., 2021; Y. Yang et al., 2019). These results suggests that the oral microbiome may be a useful tool in CRC risk assessment.

Overall, despite the work presented above in prostate, breast, and colon cancer there remains large knowledge gaps in our understanding of the oral microbiome’s applicability to population screening for these diseases. For this reason, we were interested in investigating these three cancers at a population level in both a case-control retrospective and prospective study design. To do this we leverage two different population cohorts within the Canadian Partnership Tomorrow’s Health project, the Atlantic Partnership for Tomorrow’s Health (PATH), and Alberta’s Tomorrow Project (ATP). From these two cohorts we selected saliva samples from both retrospective cases and prospective cases of prostate, breast, and colon cancer. This unique study design allowed us to investigate the relationship of these cancers with the oral microbiome both before and after diagnosis. Critically, this has allowed us to assess whether compositional changes within the oral cavity of breast, prostate, and colon cancer individuals exist both before and after diagnosis. The former being crucial for our understanding of whether the oral microbiome may be useful for the detection of individuals at risk for disease development.

## Results

### Investigation of the oral microbiome in retrospective cases of breast, prostate, and colon cancer

First, oral microbiome diversity trends between case and control samples were examined within retrospective cases of breast, prostate, and colon cancer within the Atlantic PATH cohort (Table 1). In total we examined four different alpha diversity metrics: richness, Shannon diversity, Evenness, and Faith’s Phylogenetic Diversity while controlling for DNA extraction batch. Investigation into these four metrics did not show any differences in alpha diversity between case and non-cancer controls in breast, prostate, or colon cancer (p > 0.05) (Fig. 1, Sup Fig. 1). Subsequently, we also compared three different beta diversity metrics, two that consider weighted abundances; weighted UniFrac and Bray-Curtis dissimilarity and one that considers presence/absence; unweighted UniFrac. When comparing cases of each cancer type to matched non-cancer controls, we found no significant differences in any weighted beta diversity metrics (PERMANOVA, p > 0.05) (Fig 1, Sup Fig. 2). However, we did see evidence of visual clustering in PCoA’s generated from weighted UniFrac and Bray-Curtis dissimilarity when comparing colon cancer cases to non-cancer controls (Fig 1, Sup Fig. 2C) (PERMANOVA: p=0.107, p=0.124; respectively). While comparing unweighted Unifrac distances we did find a significant difference between breast cancer cases and controls (r^2^=0.007, p=0.008), (Sup Fig 2A).

**Table 1.** Atlantic PATH cohort characteristics for retrospective cases of breast, prostate, and colon cancer.

| Cancer Type | Breast Cancer |  | Prostate Cancer |  | Colon Cancer |  |
| --- | --- | --- | --- | --- | --- | --- |
| Case vs. Control | Case | Control | Case | Control | Case | Control |
| Number of samples | 54 | 218 | 24 | 92 | 11 | 47 |
| Sex (% female) | 100% | 100% | 0% | 0% | 53% | 53% |
| Mean Age | 57.5 (8.15) | 55.6 (8.16) | 60.6 (5.92) | 57.7 (6.36) | 59.5 (10.2) | 56.9 (9.21) |
| Mean BMI | 29.0 (5.01) | 28.3 (4.72) | 29.6 (3.27) | 28.8 (3.49) | 28.9 (4.16) | 28.5 (4.13) |
| % Current Smoker | 0 | 0 | 0 | 0 | 0 | 0 |
| Median Time Since Diagnosis | 6 | N/A | 4 | N/A | 5 | N/A |

**Figure 1.**
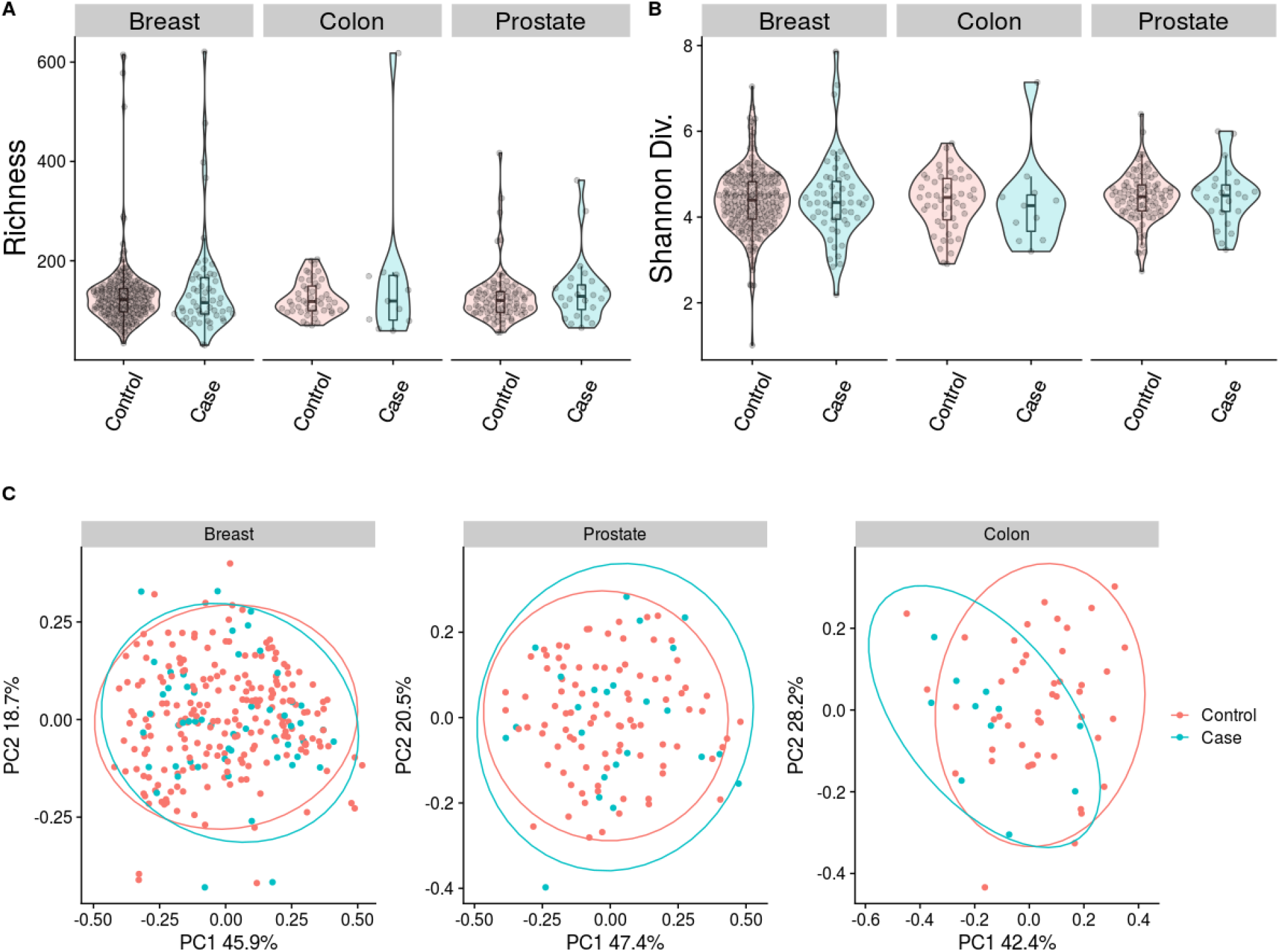
Oral microbiome diversity metrics of retrospective cases of breast, colon, and prostate cancer in the retrospective Atlantic PATH cohort. Comparing microbial diversity of non-cancer matched controls to case samples of retrospective prostate, colon and breast cancer showed no significant differences in alpha diversity as measured by richness (a), and Shannon diversity (b). Further examination into beta diversity metrics showed no significant differences in any cancer type (c) although visual clustering was identified in colon cancer (PERMANOVA; p=0.107).

To investigate whether we were missing an effect of cancer status due to the passage of time since diagnosis, we correlated each alpha diversity metric to this variable (Sup Fig. 3). We found no significant relationships except for CRC which showed a positive association between time passed since diagnosis and alpha diversity (rho=0.62, p=0.04). We also re-examined samples that were within six years of diagnosis to see whether more significant microbiome effects were present closer to cancer diagnosis. Examining these samples showed no major differences to our original analysis apart from a significant decrease in richness in CRC samples compared to matched controls (p=0.011) (Sup Fig 4).

After comprehensively examining samples for differences in diversity we decided to conduct DA analysis to identify genera or ASVs that might be associated with having previously been diagnosed with cancer. Across all cancers examined we found a total of 25 genera and 30 ASV’s associated with one or more cancer diagnoses (Fig 2, Sup Fig 5). In breast cancer we found one genus Rumminococcaecae UCG-014 that was detected as being significantly lower in relative abundance in breast cancer samples by two separate DA tools (Fig. 2). We also identified an additional 4 ASVs one of which belonged within the Rumminococcaecae UCG-014 genera (**Sup Fig 5**). The other three ASVs, two of which were classified within the Capnocytophaga genera and one within Bergeyella were all detected to be enriched in breast cancer cases (**Sup Fig 5**).

**Figure 2.**
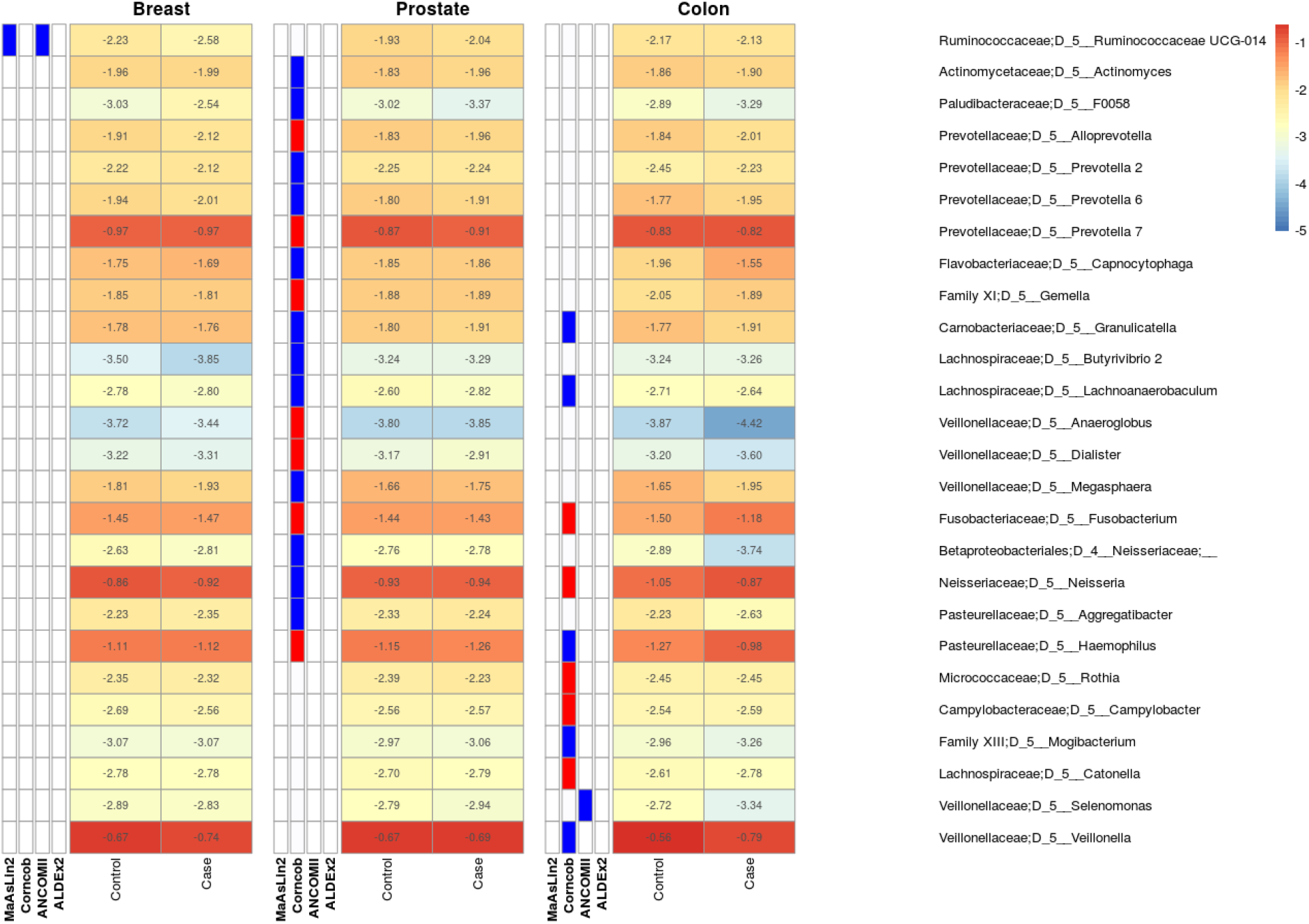
Several genera are detected as differentially abundant in the oral microbiome of retrospective cases of prostate, colon and breast cancer in the Atlantic PATH cohort. The heatmap is divided by cancer type where the first four columns represent the detection of significant associations by one of four tools: MaAsLin2, Corncob, ANCOM-II, and ALDEx2. Blue bars in the first four columns of each subgroup represent a detected increase in control samples while red bars represent a detected increase in case samples. The final two columns within each cancer sub grouping represent the log10 mean relative abundance of each genus with red representing higher abundance values and blue representing lower abundance values. Overall, corncob found the largest number of associated genera with a single genus in breast cancer also being detected by ANCOM-II.

In prostate cancer, corncob identified 19 genera and 24 ASVs that are potentially differentially abundant between case and control samples (Fig 2, Sup Fig 5). However, no other tools detected these taxa, and the overall effect size of these differences were minor ranging from 1.001 - 1.180 in log10 relative abundance mean fold changes. Furthermore, we found inconsistencies between corncob’s coefficient directionalities and the observed differences between mean relative abundances between case and control samples (Fig 2, Sup Fig 5).

In colon cancer we identified 10 genera and 6 ASVs as being differentially abundant between retrospective case and non-cancer matched control samples. All these features were identified by corncob and a single ASV classified as Fusobacterium was additionally detected by ANCOM-II to be increased in colon cancer cases (**Sup Fig. 6**). Inspection into the identity of this ASV at lower taxonomic levels using sortmeRNA (Kopylova et al., 2012) identified this ASV as potentially coming from the species *Fusobacterium periodonticum*.

Despite the relatively small taxonomic differences we found between case and control samples we were still interested in determining whether Random Forest classification models could pick up differences between case and control samples. Examining model performance on hold-out sets during cross validation showed that both breast cancer and colon cancer models performed best. Contrastingly, prostate cancer models performed at or below an AUC of 0.5 (**Fig 3A-C**). Although breast cancer models were only modestly better with AUCs ranging from 0.575 - 0.613 (**Fig 3A**).

**Figure 3:**
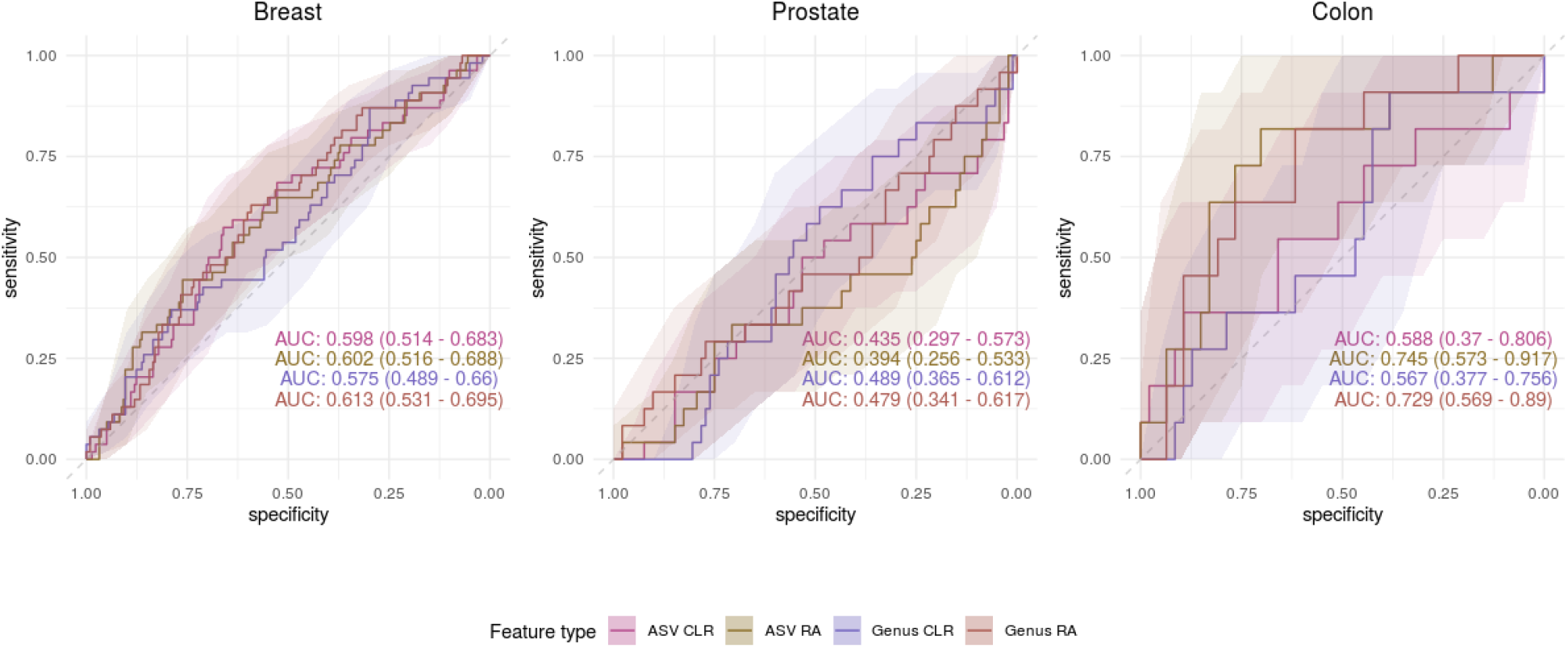
Random Forest classification of retrospective cases of breast, prostate, and colon cancer based on microbial taxonomic composition in the Atlantic PATH cohort. Receiver operator curves (ROC) showing the specificity and sensitivity of the classification of non-cancer matched controls and retrospective cases of breast, prostate or colon cancer. Models were constructed using 100-repeat 5-fold cross validation and hold-out performance was determined through taking the mean number of votes for each hold-out sample across all 100 repeats. Within each plot four different ROCs are represented showing the classification accuracy using ASVs or genera normalized with either total-sum-scaling or center-log-ratio abundance. Shaded areas represent 95% confidence intervals determined through 2000 bootstrap samplings.

Colon cancer models performed the best although large confidence intervals were observed due to low sample size. Furthermore, colon cancer models showed highly variable performance depending on feature normalization. Models built using center-log-ratio normalizations performed only slightly better than random expectation with ASVs having an AUC of 0.588 (0.370 - 0.806 95% CI), and genera having an AUC of 0.567 (0.377 - 0.756 95% CI). However, models built using relative abundances showed stronger results with models having AUCs ranging from 0.729 - 0.745 (**Fig 3**). Examining the top 10 most important features of each of these models showed relatively small decreases in accuracy for any single ASV/genera (**Sup Fig 7-8**). Indeed, further inspection also showed that of these ASV/genera only 1 ASV overlapped with the previously identified differentially abundant taxon (**Sup Fig. 6, Sup Fig 7**). This ASV was classified into the genera Veillonella and upon further inspection, best aligned to *Veillonella atypica* (100% identity) within the SILVA V138 database.

### Investigation of the oral microbiome in prospective cases of breast, prostate and colon cancer

We next decided to investigate if compositional changes within the oral microbiome are present before the diagnosis of breast, prostate, and colon cancer. Like our retrospective analysis we first examined changes in overall microbial community structure by looking for differences in alpha and beta diversity between cancer cases and matched non-cancer controls. We found no significant differences in alpha diversity in either cohort using linear models comparing case vs. control in any of the four metrics (**Fig 4, Sup Fig 9-10**). Correspondingly we did not find any significant differences in beta diversity between breast, colon, or prostate cancer cases and matched non-cancer controls in weighted UniFrac, unweighted UniFrac, or Bray-Curtis dissimilarity (**Fig 4. Sup Fig 11-12**) (PERMANOVA p > 0.05).

**Figure 4:**
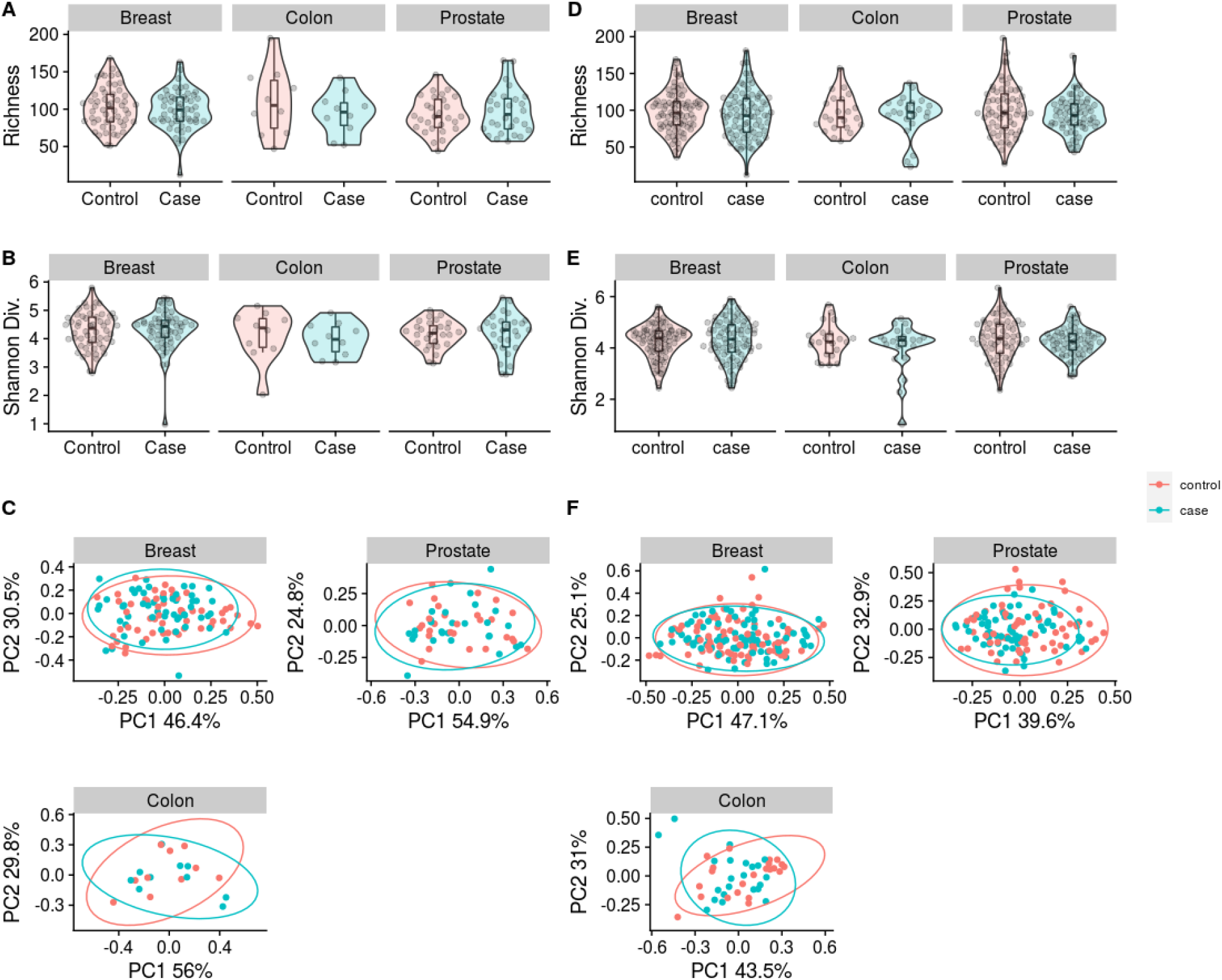
Oral microbiome diversity metrics of prospective cases of breast, prostate and colon cancer in Atlantic PATH and ATP cohorts. Oral microbiome diversity analysis comparing non-cancer matched controls to prospective cases of colon breast and prostate cancer. Alpha diversity analysis as measured by richness (A) and Shannon diversity (B) as well as beta diversity measured as weighted UniFrac (C) showed no significant differences in the prospective PATH cohort. Similarly no significant differences in richness (D), Shannon diversity (E) or weighted UniFrac (F) in any cancer type in the ATP dataset.

Like our previous retrospective analysis, we also examined whether the time between sample collection and diagnosis had a major impact on signal within the case samples. Spearman correlations showed no significant relationships in any of the alpha diversity metrics in both cohorts (**Sup Fig 13-14**) (p > 0.05). Furthermore, examining case samples that were collected within four years of diagnosis showed similar results except for prostate cancer in the Atlantic PATH cohort which showed a significant increase in Faith’s phylogenetic diversity (p=0.025) and richness (p=0.041) in case participants (**Sup Fig 15-16**).

After examining overall oral microbial community structure through various diversity metrics we were interested in determining whether there was any evidence of specific ASVs or genera being associated with disease status. In both prospective cohorts (Atlantic PATH, ATP) we found no genera being associated with disease status, however, we did find a small number of ASVs associated with disease status in both cohorts. In the Atlantic PATH cohort, we found an increase in the relative abundance of an ASV classified as Alloprevotella rava in prostate cancer (**Sup Fig 17-18**). We additionally found a decrease in the relative abundance of an ASV classified as Streptococcus in colon cancer (**Sup Fig 17-18**). Interestingly, none of these ASVs overlapped with those identified in the ATP dataset. Within the ATP cohort we detected two ASVs being decreased in relative abundance in colon cancer, although the significance of these taxa was mostly driven by outliers within control samples (**Sup Fig 19-20**). Within this cohort ANCOM-II also detected that the relative abundance of an ASV classified to an uncultured Stomatobaculum to be decreased in prospective breast cancer samples (**Sup Fig 19-20**).

As with our previous analysis we were interested in applying Random Forest models to each prospective cancer type to help identify whether disease signatures exist within the oral microbiome. Separate models were generated for each cancer type and cohort. Overall, models for breast cancer performed poorly in both cohorts with accuracies similar to random classification in both Atlantic PATH and ATP cohorts (**Sup Fig 21-22**). Interestingly, in the case of prostate cancer three of four models in the Atlantic PATH cohort performed slightly above random classification with AUCs ranging from 0.602 - 0.665 (**Sup Fig 22**). However, in the ATP cohort all models performed at or below an AUC of 0.5 (**Sup Fig 23**) although it should be noted that 95% confidence intervals on these AUCs were large due to small sample sizes (**Table 2-3**).

**Table 2.** Atlantic PATH cohort characteristics for the investigation of the oral microbiome in prospective cases of breast, colon, and prostate cancer.

| Cancer Type | Breast Cancer |  | Prostate Cancer |  | Colon Cancer |  |
| --- | --- | --- | --- | --- | --- | --- |
| Case vs. Control | Case | Control | Case | Control | Case | Control |
| Number of samples | 54 | 54 | 28 | 28 | 10 | 10 |
| Sex | 100% |  | 0% |  | 70% |  |
| Mean Age | 56.6<br>(7.74) | 56.9<br>(8.38) | 60.6<br>(4.43) | 61.0<br>(4.93) | 60.4<br>(7.88) | 60.7<br>(8.25) |
| Mean BMI | 26.9<br>(5.25) | 26.9<br>(5.45) | 28.3<br>(4.46) | 28.1<br>(4.72) | 27.9<br>(6.08) | 28.3<br>(5.80) |
| % Current Smoker | 1.85% | 3.70% | 0% | 3.57% | 0% | 0% |
| Median Time Before<br>Diagnosis | 4 | N/A | 3.5 | N/A | 3 | N/A |

**Table 3.** ATP cohort characteristics for investigation of the oral microbiome in prospective cases of breast, colon, and prostate cancer.

| Cancer Type | Breast Cancer |  | Prostate Cancer |  | Colon Cancer |  |
| --- | --- | --- | --- | --- | --- | --- |
| Case vs. Control | Case | Control | Case | Control | Case | Control |
| Sex (% Female) | 100% |  | 0% |  | 50% |  |
| Number of<br>samples | 82 | 82 | 64 | 64 | 22 | 22 |
| Mean Age | 57.7<br>(8.74) | 57.7 (8.70) | 63.2 (6.92) | 63.2 (6.88) | 60.0 (10.0) | 60.0 (9.99) |
| Mean BMI | 28.3<br>(6.00)<br>11 N/a | 27.1 (5.70)<br>9 N/a | 27.3 (4.3)<br>7 N/a | 28.2 (4.93)<br>5 N/a | 30.6 (6.94)<br>1 N/a | 27.1 (4.49)<br>3 N/a |
| Current Smoker | 1.22% | 1.22% | 15.6% | 15.6% | 9.09% | 9.09% |
| Median Time<br>Before Diagnosis | 4.28 | N/A | 3.10 | N/A | 2.98 | N/A |

Finally, models of prospective cases of colon cancer showed variable results between the two cohorts of interest. With models in the Atlantic PATH cohort showing low performance AUCs ranging from 0.380 - 0.620 (**Fig 5, Sup Fig 21**). Contrastingly, in the ATP dataset stronger classification accuracies were found when using center-log-ratio normalizations with the genera level model performing the best (AUC 0.717; 95% CI: 0.549 - 0.884) (**Fig 5, Sup Fig 22**).

**Figure 5:**
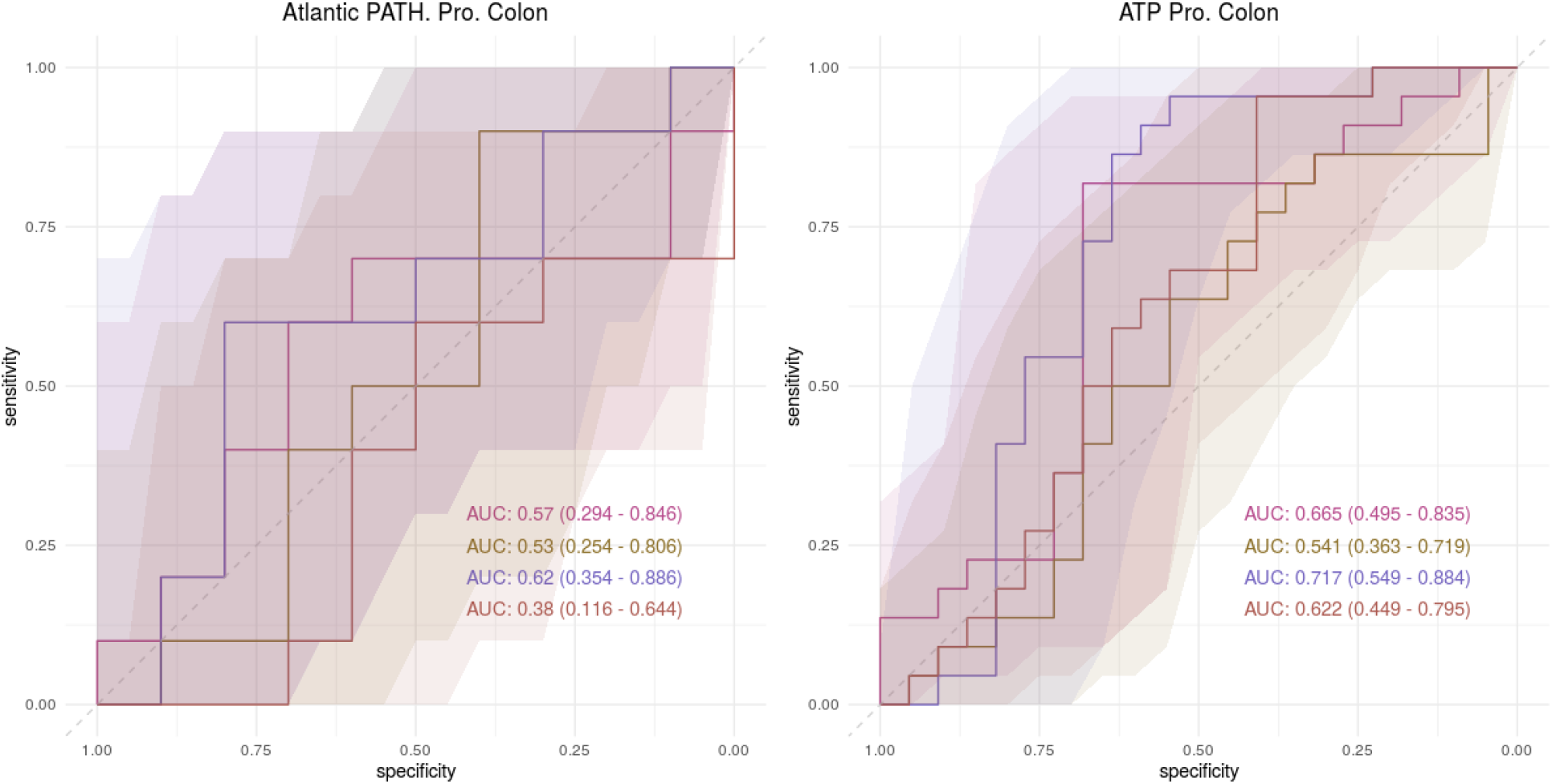
Random Forest Classification performance of prospective cases of colon cancer in the Atlantic PATH and ATP cohorts. Receiver operator curves (ROC) showing the specificity and sensitivity of the classification of non-cancer matched controls and prospective cases of colon cancer in the PATH and ATP datasets. Models were constructed using 100-repeat 5-fold cross validation and hold-out performance was determined through taking the mean number of votes for each hold-out sample across all 100 repeats. Within each plot four different ROCs are represented, showing the classification accuracy using ASVs or genus normalized with either total-sum-scaling or center-log-ratio abundance. Shaded areas represent 95% confidence intervals determined through 2000 bootstrap samplings.

Inspecting the top ten most important genera through out-of-bag permutation analysis within our CLR normalized colon cancer model showed no single genera as being particularly important to classification accuracy (accuracy decrease ranging from: 0.003 - 0.016). The most important genera within the model only decreased out-of-bag accuracy by 0.016 (SD; 0.002) although inspection of its CLR abundance did show a notable increase in case samples when compared to non-cancer controls. Inspection of other important genera within this model showed interesting CLR abundance patterns although as mentioned previously none were identified in our previous differential abundance analysis (**Fig 6**).

**Figure 6:**
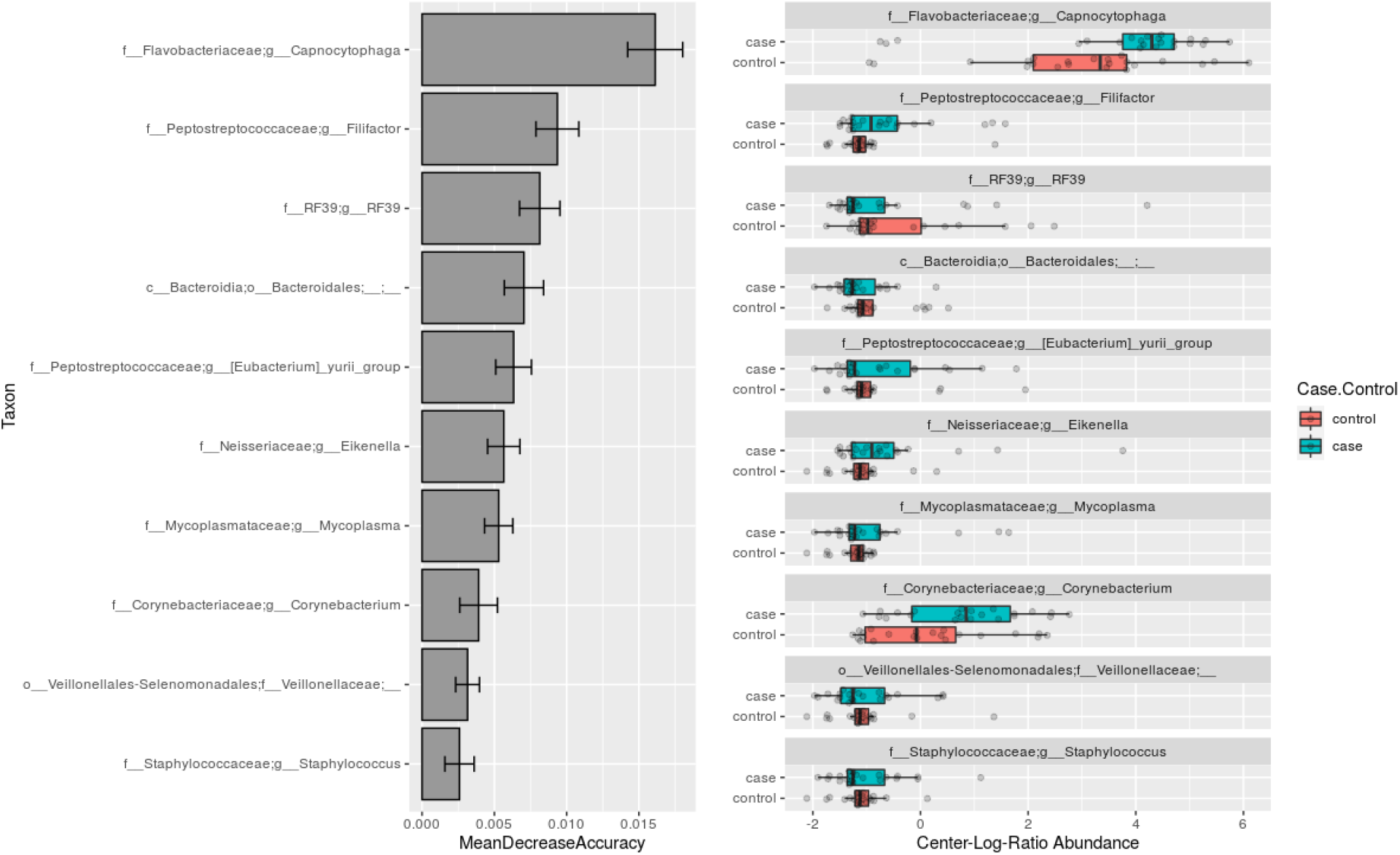
Feature importance of genus level center-log-ratio normalized Random Forest classification of prospective cases of colon cancer in the ATP dataset. Feature importance was determined using out-of-bag permutation feature analysis. MeanDecreaseAccuracy represents the mean out-of-bag accuracy loss when that feature was randomly permuted across samples (A). Feature center-log-ratio abundance patterns are shown in panel (B) and show possible examples of interesting genera to further investigate in future studies.

## Discussion

Herein we examined the oral microbiome in the context of both retrospective and prospective cases of prostate, colon, and breast cancer in a population setting. Our analysis showed no significant changes in oral microbiome diversity in either retrospective or prospective cases of these cancers. Although we did find evidence of visual clustering of retrospective colon cancer cases when examining weighted beta diversity metrics. Investigating the relationships of individual taxon and cancer status showed some evidence of potential associations, although the majority were only detected by one of four DA tools indicating a low level of evidence. Accordingly, Random Forest classification of case samples and non-cancer matched controls showed relatively low classification accuracies with colon cancer showing the strongest signal in both retrospective and prospective analysis. Overall, our findings suggest that no large community changes exist in the oral microbiome of individuals with retrospective or prospective cases of prostate and breast cancer. Although a minor amount of evidence in our report does suggest there may be potential individual taxon relationships within these diseases. Contrastingly, through Random Forest modeling we did find some signal in retrospective cases of colon cancer and in one of our prospective colon cancer cohorts. Highlighting that future studies on prospective colon cancer cases are warranted.

Examining our results in breast cancer more closely showed strong concordance with previous work by Wang et al., who also found no changes in overall oral microbiome composition in United States individuals with breast cancer (H. Wang et al., 2017). These results contrast with a recent study by Wu et al., who identified differences in microbial diversity and the abundance of Porphyromonas and Fusobacterium (Wu et al., 2022). This could be due to several reasons including the fact that these studies were conducting under highly different populations, as geographic differences have been shown to impact oral microbiome composition (Li et al., 2014). Unlike either of these studies, we did find evidence for a modest decrease in the relative abundance of Ruminococcaceae UCG-014 an uncultured genus that we previously linked to differences in height within healthy individuals of the same cohort (Nearing et al., 2020). Whether this taxon plays any role in disease status is unclear, however, due to this association not being recovered in either of our prospective cohorts it’s likely that its association is limited to during or soon after cancer development. Similarly, we also identified an increase in two ASVs classified within the genus *Capnocytophaga* which were not detected in our prospective cohorts. Interestingly, within this genus, *C. gingivalis* has previously been associated with oral squamous cell carcinomas (Healy & Moran, 2019). Moreover, recent work has shown that supernatant from this species has the potential to enhance cellular invasion and migration of tumor cells (W. Zhu et al., 2022). Highlighting that this species may play a role in disease development or progression.

Investigating signal within the oral microbiome of breast cancer individuals using Random Forest modeling showed relatively little signal with accuracies in retrospective cases only slightly better than random assignment. Based on these results, we believe it is unlikely that the oral microbiome could be used as a biomarker to detect the risk of breast cancer development within a population setting.

To the best of our knowledge this report is the first to examine the relationship of the oral microbiome and prostate cancer diagnosis. Similar, to breast cancer we found no large shifts in oral microbiome diversity in prospective or retrospective cases of prostate cancer. Although, we did see a possible time dependent effect in the Atlantic PATH cohort which was not found in our second ATP cohort. Whether these differing results are due to DNA extraction, regional differences, or simply a false positive discovery would require further investigation in future studies.

Despite not identifying any consistent differences in diversity, multiple ASVs and genera were identified by corncob to be associated with retrospective cases of disease. Unsurprisingly, comparing these results to those previously identified within the gut showed little overlap (Golombos et al., 2018; Liss et al., 2018; Matsushita et al., 2021). Additionally, none of these retrospective taxonomic relationships were recovered in our prospective datasets. These results suggest that these retrospective associations may be related to other broad lifestyle changes that occur after prostate cancer diagnosis. Indeed, several of these retrospective taxa associations were previously associated with various lifestyle, dietary and anthropometric measurements within healthy PATH participants (Nearing et al., 2020). This highlights that many microbes, even those potentially associated with disease, are often affected by multiple daily life factors some of which could be associated with disease diagnosis.

Accompanying these results, we saw little to no signal in our Random Forest prostate cancer classification models. Some signal was recovered from the best models trained on the Atlantic PATH cohort (AUROC 0.665); however, due to small sample sies (N=28) confidence intervals remained large. Moreover, this signal was not recovered in our additional ATP cohort. These results suggest that the oral microbiome is unlikely to be a strong biomarker of prostate cancer risk.

Colon cancer showed the strongest evidence for differences in oral microbiome diversity, though, none of these differences were detected as being significant. Despite this we did see evidence of visual clustering in weighted UniFrac profiles of retrospective cases of disease (p=0.102, p=0.124). This matches with previous work by both Flemer et al., and Wang et al., who found significant differences in oral microbiome beta diversity between healthy controls and individuals diagnosed with colon cancer (Flemer et al., 2017; Y. Wang et al., 2021). Further investigation of diversity metrics also showed a reduction in oral microbiome richness in individuals that were diagnosed within 6 years of sample collection. Interestingly, this conflicts with previous reports by Wang et al., who found increases in oral microbiome diversity within CRC patients (Y. Wang et al., 2021). However, it should be noted that this could be due to several different factors including those associated with differing treatment regimens, sample timing, or environmental exposures.

In retrospective cases of colon cancer, we found evidence of an increase in an ASV classified as *Fusobacterium peridonticum* by two different microbiome DA frameworks (corncob, ANCOM-II). The relative abundance of *Fusobacterium peridonticum* a close relative to *Fusobacterium nucleatum* has previously been identified as being increased in the oral microbiome of oral small cell carcinoma (C.-Y. Yang et al., 2018), head and neck cancer (Mougeot et al., 2021), and pancreatic cancer patients (Sun et al., 2020). Accordingly, whether the relative abundance of this oral microbe represents a specific connection to colon cancer is still in question as it may represent a broader connection to other events associated with cancer diagnosis. Finally, of the three cancers examined, colon cancer showed the most consistent signal in our Random Forest modeling, although substantial differences in classification accuracies were noted between our two prospective cohorts. This could have been due to a few factors including sample size differences, collection method, or possibly differences in health risk factors between Atlantic Canada and Alberta. Unfortunately, attempts to train Random Forest models on combined datasets were not successful due to different collection and DNA extraction approaches causing significant bias between studies (Nearing et al., 2021; Pollock et al., 2018).

One puzzling result we noticed from our colon cancer Random Forest models, was that the type of normalization used had large impacts on model accuracies and was not consistent between retrospective and prospective datasets. We found total-sum-scaling to perform the best in our retrospective cohort but found center-log-ratio transformation to perform better in our prospective ATP cohort. Whether this has biological significance is unclear and suggests that future models may be interested in testing several different data normalizations during model training.

Within our analysis we have also identified several limitations that should be noted when reviewing our results. The first is that in the case of both prostate and colon cancer our sample size numbers are small in all three datasets we examined (Table 1-3). These smaller samples most likely interfered with our ability to detect small differences in both community composition and individual taxa abundances. A second limitation of our study is the different extraction methods used for samples that came from the two different population cohorts, Atlantic PATH and ATP. This along with other technical variations such as differences in sample collection led to the need to conduct stratified analysis reducing our ability to valid signal between cohorts. Finally, we would like to acknowledge that our datasets were relatively homogenous, with our dataset being predominantly from white Canadians, with income and education levels above average Canadian census data (Sweeney et al., 2017; Ye et al., 2017). As such this significantly limits our ability to identify distinct oral microbial signatures in groups that are disproportionately affected by cancer development.

In conclusion, we believe that our report shows that the oral microbiome is unlikely to be an effective population-based risk marker for cases of prostate or breast cancer, although changes in specific bacterial abundances within these diseases may still exist. Contrastingly, in the case of colon cancer our work indicates that disease status is related to changes in the oral microbiome and may be useful as a risk marker for colon cancer development. Future studies should aim to evaluate when oral microbiome changes occur in prospective colon cancer cases to determine its suitability for risk stratification.

## Methods

### Study design

This report includes the analysis of saliva samples from individuals who had previously been enrolled in two regional cohorts within the Canadian Partnership for Tomorrow’s Health project, a pan-Canadian prospective cohort study focused on examining the influence of genetics, the environment, and lifestyle factors on Canadian’s health. The regional cohorts of interest for this study include Atlantic PATH (which includes participants from the 4 Atlantic provinces: (Nova Scotia, New Brunswick, Prince Edward Island, and Newfoundland and Labrador) and ATP (participants from the western province of Alberta). For this study, both retrospective (cases diagnosed prior to baseline data and sample collection) and prospective (cases diagnosed after baseline data and sample collection) nested case-control designs were employed. This study was granted ethics approval from Dalhousie University Health Sciences Research Ethics Board (REB #2018-4420).

### Atlantic Partnership for Tomorrow’s Health cohort characteristics

At baseline, demographics, lifestyle, personal and family medical history were self-reported on questionnaires, and a subset attended assessment centers where physical measurements and biospecimens such as saliva were collected. For more details on baseline characteristics of the Atlantic PATH cohort, an in depth descriptive cohort profile has been previously published (Sweeney et al., 2017). Follow-up questionnaire data was collected between 2016-2019.

For the purposes of this study sample selection within the Atlantic cohort was divided into either a retrospective or prospective nested case-control design as previously described. Prior cancer diagnosis was determined through baseline questionnaires filled out by each participant. All available breast, prostate, and colon cancer case saliva samples were included in this study, and control samples (non-cancer) were selected to match cases (1:5) by sex, age (+/-3 years), BMI (+/-3), and smoking status (current vs never/former). The retrospective design included 588 saliva samples from the Atlantic PATH biospecimen repository based on case and non-cancer control matches to individuals that had been diagnosed with breast (n=61), prostate (n= 23), or colon cancer (n= 14) prior to baseline saliva collection.

For the prospective design, new incident cases of cancer were determined through follow up questionnaire surveys filled out by each participant. A one-to-one case control design was used with non-cancer control samples being matched to case samples based on age (+/- 3 years), sex, and BMI (+/- 3). A minor number of current smokers ranging from 0% - 3.70% depending on cancer status were included in this analysis (Table 2). The prospective design included 230 samples from the Atlantic PATH cohort who had breast (n=67), prostate (n=35), or colon cancer (n=13).

The median length of time between sample collection and cancer diagnosis for each cancer can be found in Table 1 (retrospective) and Table 2 (prospective) along with other sample characteristics broken down by cancer type, study design, and case control status.

### Alberta’s Tomorrow Project cohort characteristics

Recruitment and baseline data collection took place between 2000 and 2015 with biospecimen collection beginning in 2009. Details on cohort characteristics, recruitment, and design have been previously published (Ye et al., 2017).

ATP collected self-reported baseline and follow-up questionnaire data on demographics and health risks. New incident cases of cancer were confirmed through linkage to Alberta Cancer Register (ACR). Case and control samples were matched in 1:1 design based on age (+/- 2 years), sex, and smoking status (current, former, never). In total 414 saliva samples were identified from ATP’s biospecimen repository based on non-cancer controls and cases of breast (n=102), prostate (n=76), or colon cancer (n=29) that were diagnosed after saliva sample collection. The median length of time between saliva collection and cancer diagnosis along with other relative metadata can be found in Table 3.

### Oral microbiome 16S rRNA gene sequencing

Samples from the Atlantic PATH cohort were processed as previously described (Nearing et al., 2020). Frozen saliva samples were stored at -80C and then thawed at room temperature and aliquoted into 96 well plates. In a biosafety cabinet using standard sterile techniques DNA was extracted using a QIAamp 96 PowerFecal QIAcube HT kit following the manufacturer’s instructions using a TissueLyser II and the addition of Proteinase K at the indicated optional step. Sequencing was done at the Integrated Microbiome Resource at Dalhousie University. PCR amplification of the V4-V5 16S rRNA gene region was done using V4-V5 fuson primers (515FB - 926R) and a high-fidelity Phusion polymerase. A total of 25 cycles of PCR were done: denaturing at 98℃, annealing at 55℃, and elongating at 72℃. Sequencing was then conducted using an Illumina MiSeq to produce 300-bp demultiplexed paired-end reads.

Samples from the Alberta’s Tomorrow Project cohort were collected using an Oragene® DNA OG-250 kit manufactured by DNA Genotek. Samples were collected either in person at local study centers or saliva sample kits were sent to participants by regular postal mail with a return envelope included. Participants were instructed not to eat, chew gum, or smoke 30 minutes prior to providing a saliva sample. They were asked to spit into the container until the saliva reached the indicated level, screw the cap on, shake for 10 seconds and send the sample back through the mail. DNA from samples were then extracted using a DNA Genotek PrepIT PT-LP2 kit. After extraction samples were sequenced at the Integrated Microbiome Resource in the same manner as samples from the Atlantic PATH cohort.

### 16S rRNA gene sequence processing

Processing of 16S sequencing data was conducted as previously described (Nearing et al., 2020). Primers were removed using cutadapt with default settings (M. Martin, 2011). Primer-free reads were stitched using the QIIME2 VSEARCH join-pairs plugin (Bolyen et al., 2019; Rognes et al., 2016). Stitched reads were then filtered with default settings using the QIIME2 plugin q-score-joined. Reads were then corrected into amplicon sequence variants using the QIIME2 plugin Deblur with a trim length of 360 bp, and one read set as the minimum number required to pass filtering (Amir et al., 2017). For each dataset examined, 0.1% of the mean sample depth was calculated and ASVs below this abundance across all samples within that dataset were removed. This is to keep in line with the previously described Illumina MiSeq bleed-through rate. ASVs were placed into the Greengenes 13_8 99% reference 16S rRNA tree using the QIIME2 fragment-insertion SEPP plugin (DeSantis et al., 2006; Janssen et al., 2018; MIRARAB et al., 2011). Rarefaction curves were generated for each dataset separately and a suitable rarefaction depth of 5,000 was chosen for the Atlantic PATH retrospective cohort and 3,000 for the Atlantic PATH prospective and ATP prospective cohorts. Rarefied data was used to generate both alpha and beta diversity metrics. Samples below these sequencing depths along with those that had no remaining case or control samples were removed from further analysis. Additionally, a single sample in the ATP prospective dataset was removed due to significant contamination during sample preparation. Final case-control sample numbers for each dataset that pass all quality filtering are presented in Tables 1-3. ASVs were then assigned taxonomy using a naive Bayesian QIIME2 classifier trained on the 99% Silva V138 16S rRNA database (Bokulich et al., 2018; Quast et al., 2013).

### Microbial Diversity Analysis

Alpha and beta diversity metrics were generated using the QIIME2 command “core-metrics-phylogenetic” with the previously described rarefaction values and phylogenetic tree. Diversity matrices were then exported into R and analyzed between case and control samples for each separate cohort and study design. Alpha diversity between case/control samples were examined using linear models with the inclusion of an “extraction_run” covariate for the Atlantic PATH retrospective samples due to the large number of batch extractions and amount of time passed between sample extractions. In total four different alpha diversity metrics were investigated: Faith’s phylogenetic diversity, Shannon diversity, evenness, and richness. An alpha value of 0.05 was chosen as our significance threshold before conducting any statistical analysis. Violin boxplots were generated using ggplot2 while jitter points were added using the R package ggbeeswarm (Wickham, 2009).

Beta diversity metrics were compared using a PERMANOVA test between case samples and case matched control samples for each cancer type within each cohort and study design using the ‘adonis2’ function within the vegan R package (Dixon, 2003). In the case of the Atlantic PATH retrospective data we included the covariate extraction number due to the large number of different extraction runs and time taken between sample extractions for this dataset. An alpha value of 0.05 was chosen before any statistical testing was conducted. A secondary PERMANOVA analysis was also conducted between all cancer types within each cohort and study design followed by pairwise tests between cancer type and all controls within that cohort and study design. In total three different beta diversity metrics were examined: weighted UniFrac, unweighted UniFrac and Bray-Curtis dissimilarity. These three beta diversity metrics were visualized using principal coordinate analysis using the function cmdscale within an R programming environment and ggplot2. Ellipses were added to each sample type using the function ‘stat_ellipse()’.

### Microbial Differential Abundance Analysis

Differential abundance analysis was conducted using four different tools developed to analyze microbiome data: ALDEx2 (Fernandes et al., 2014), ANCOM-II (Kaul et al., 2017; Mandal et al., 2015), corncob (B. D. Martin et al., 2020), and MaAsLin2 (Mallick et al., 2021). These tools range in their consistency and power to detect differences between groupings and should give a broad range on the ability to detect differentially abundant taxa (Nearing et al., 2022). Each tool was run at both the ASV and genus taxonomic levels. All tools were run comparing taxonomic abundance against case versus control status. Each tool was run separately for each cancer type, cohort, and study design. During the examination of the Atlantic PATH retrospective dataset we also included DNA extraction as a covariate due to not all samples being extracted at the same time. For all tools, taxa that were not found in at least 5% of samples were removed from consideration. Filtered p-values were then corrected for false discovery using Benjamini– Hochberg correction (Benjamini & Hochberg, 1995).

ALDeX2 analysis was run using default settings and general linear models. This includes using a center-log-ratio transformation, and 128 Monte Carlo samplings to generate probability distributions from the observed count data.

ANCOM-II was run using scripts available at: https://github.com/FrederickHuangLin/ANCOM-Code-Archive. Genus and ASV abundance tables were first processed using the function “feature_table_pre_process”. The main grouping variable of interest during pre-processing and the determination of structural zeros was case versus controls. A value of 0.05 was used to determine outlier zeros and outlier values. Pre-processed tables were then passed into the main ANCOM function with the inclusion of DNA extraction batch as a covariate when examining retrospective PATH data. Significance was determined using a percentage cutoff of 70% for the w statistic.

Corncob was run by first importing taxonomic abundance tables and their corresponding metadata into phyloseq objects (McMurdie & Holmes, 2013). The function differentialTest was then run using the above phyloseq object with the “wald” test option. The phi formula was set to match the phi-null formula to control for differences in variability across sample groupings.

MaAsLin2 was run using default settings and an arcsine transformation. Case versus control was used as a fixed effect. In the case of the PATH retrospective dataset an additional fixed effect of the DNA extraction batch was included.

### Random Forest model training

Random Forest models were trained and used to classify case and control samples from each dataset and study design. Training and classification were done using 100-repeat-5-fold cross validation. In the retrospective dataset control samples were randomly downsampled within each fold training session to avoid unbalanced model training and biasing data within the hold-out fold. In all datasets taxon found in less than 5% prevalence were filtered out prior to model training. After training and cross validation, the mean number of votes on each hold-out set across all repeats was then calculated. Receiver operator curves were constructed using pROC and confidence intervals were estimated using 2000 bootstrap replicates (Robin et al., 2011). Variable importance was calculated from models trained on the entire dataset by determining the difference in the out-of-bag prediction error rate after the variable of interest was permuted.

## Data availability

All sequence data has been uploaded to the European Nucleotide Archive and are available under the accession numbers PRJEB38175 and PRJEB56605. Code to analyze all data is available on GitHub at https://github.com/nearinj/Oral_Microbiome_Prostate_Breast_Colon_Cancer. A subset of deidentified metadata used in this project can also be found at the above GitHub link. Additional metadata variables can be accessed by contacting either the Atlantic Partnership for Tomorrow’s Health project or the Alberta’s Tomorrow Project.

## Acknowledgements

This research was conducted using data and biosamples from both Atlantic PATH and Alberta’s tomorrow project under the application numbers 2018-103 and 2020-06-Langille respectively. We would like to thank all participants in both studies who donated their time, personal medical history, and saliva samples to this project. We would also like to thank both the Atlantic PATH and Alberta’s tomorrow project team members for assistance during research applications, data collection, and management. J.T.N. is supported by both a Research Nova Scotia, Scotia Scholars award (2019 - 2022) and a Nova Scotia Graduate Student Scholarship (2019-2023). The data used in this research was made available by both the Atlantic Partnership for Tomorrow’s Health and Alberta’s Tomorrow Project, which are regional cohorts of the Canadian Partnership for Tomorrow’s Health Project, funded by the Canadian Partnership Against Cancer, Health Canada, Alberta Health, and the Alberta Cancer Foundation. Cancer registry data was obtained through linkage with Surveillance & Reporting, Cancer Research & Analytics, Cancer Care Alberta. The views expressed here represent the views of the authors and do not necessarily represent the views of Atlantic PATH, Alberta Tomorrow Project, or any of its funders.

## Author contributions

JTN, VD, and MGIL proposed the project; VD and MGIL supervised the project; JTN processed and prepared datasets; analyzed datasets; generated figures; and wrote the paper. VD and MGIL provided constructive feedback during manuscript preparation. All authors approved the final manuscript.

