## Supplemental Information for "Investigating the oral microbiome in retrospective and prospective cases of prostate, colon, and breast cancer"

### Supplemental Figures

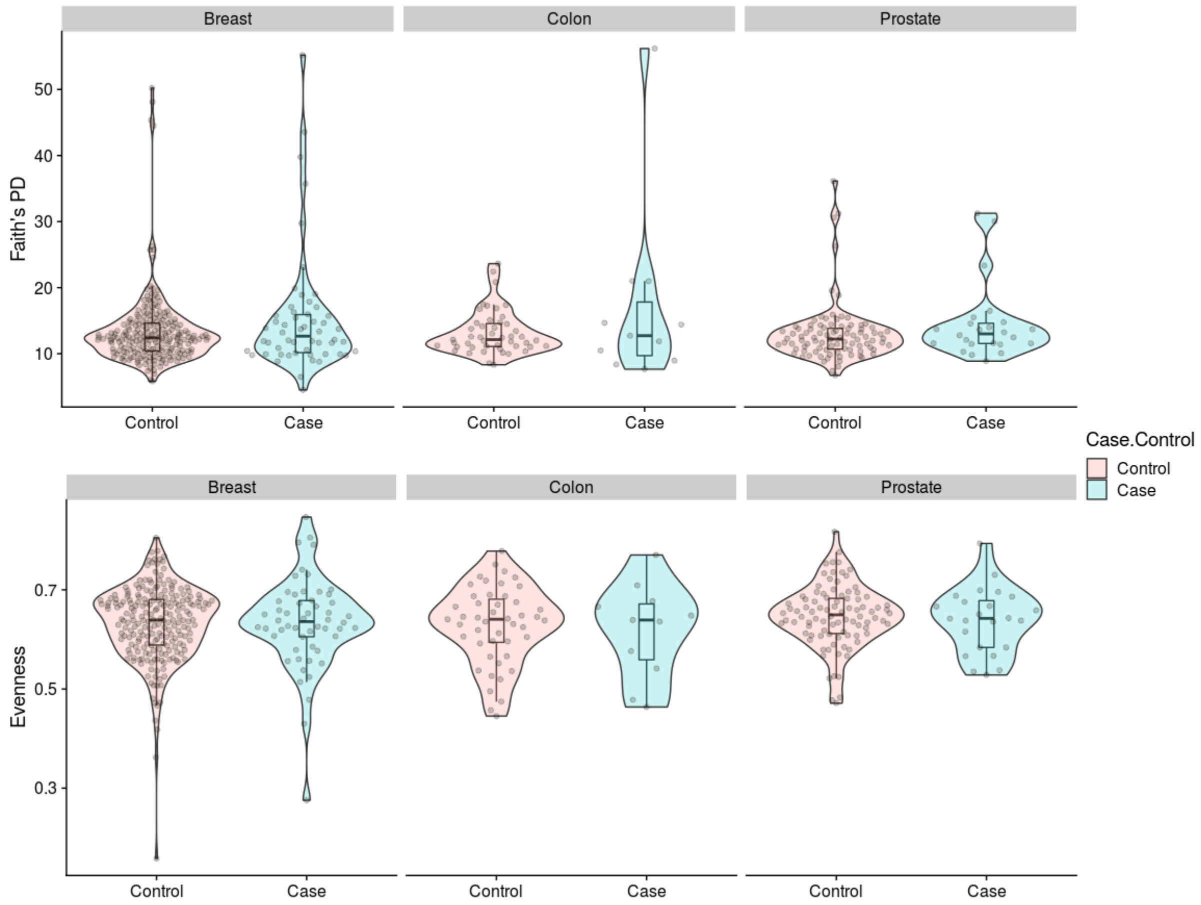


**Supplemental Figure 1. Alpha diversity metrics comparing retrospective cases of breast, prostate, and colon cancer to matched controls in the Atlantic PATH cohort.** Faith's phylogenetic diversity (top row) and evenness (bottom row) between non-cancer matched controls and retrospective cases of breast, colon, or prostate cancer. We found no significant differences ($p>0.05$) using linear models while controlling for DNA extraction. Boxplots represent the median and interquartile while violin plots represent the density.

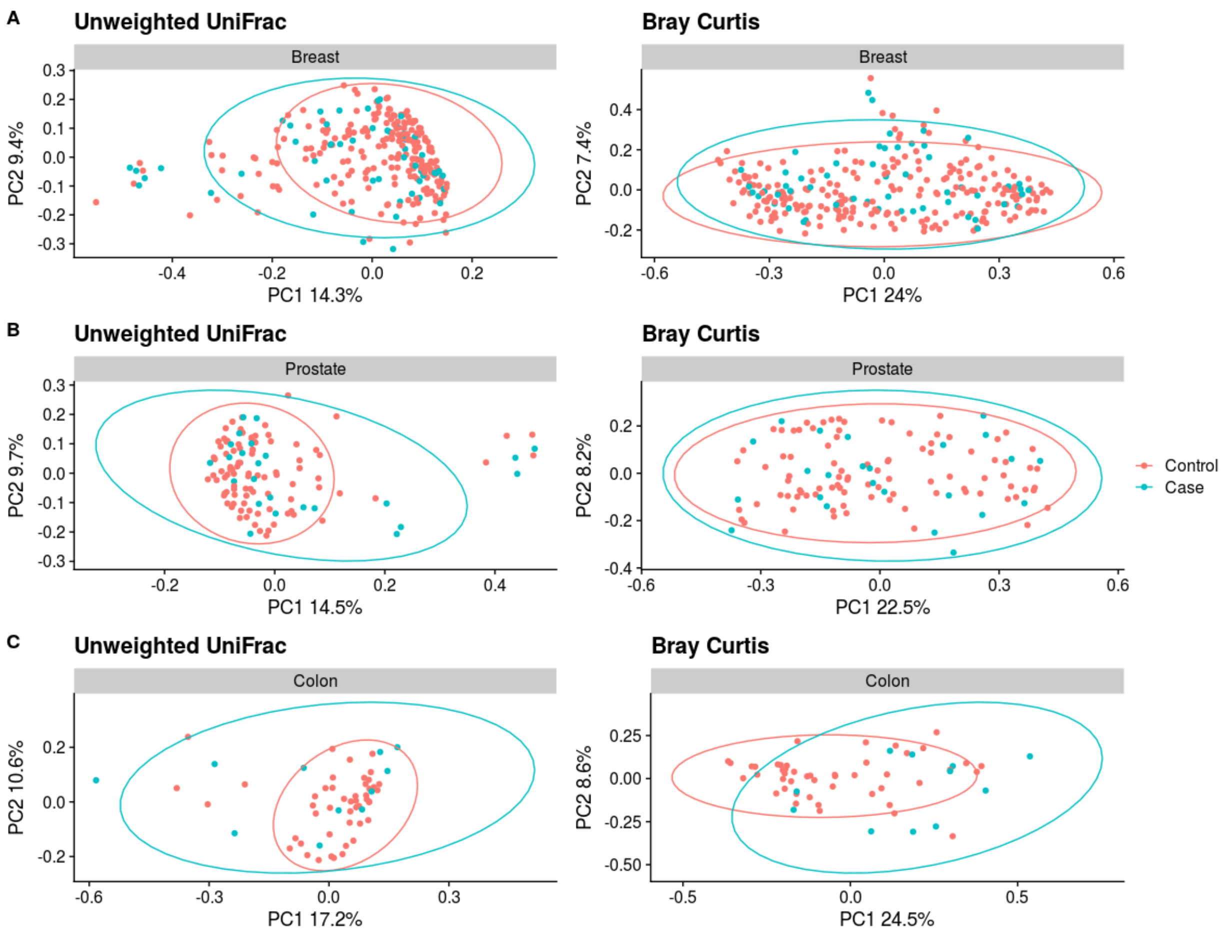


**Supplemental Figure 2. Case-Control beta diversity analysis of each cancer type within the retrospective Atlantic PATH cohort.** Comparison of two different beta diversity metrics (unweighted UniFrac and Bray-Curtis dissimilarity) between non-cancer matched controls and retrospective cases of breast (A) prostate (B) and colon cancer (C). Using PERMANOVA tests while controlling for extraction we found a significant difference in unweighted UniFrac distances in breast cancer case samples ($r^2$=0.007, p=008).

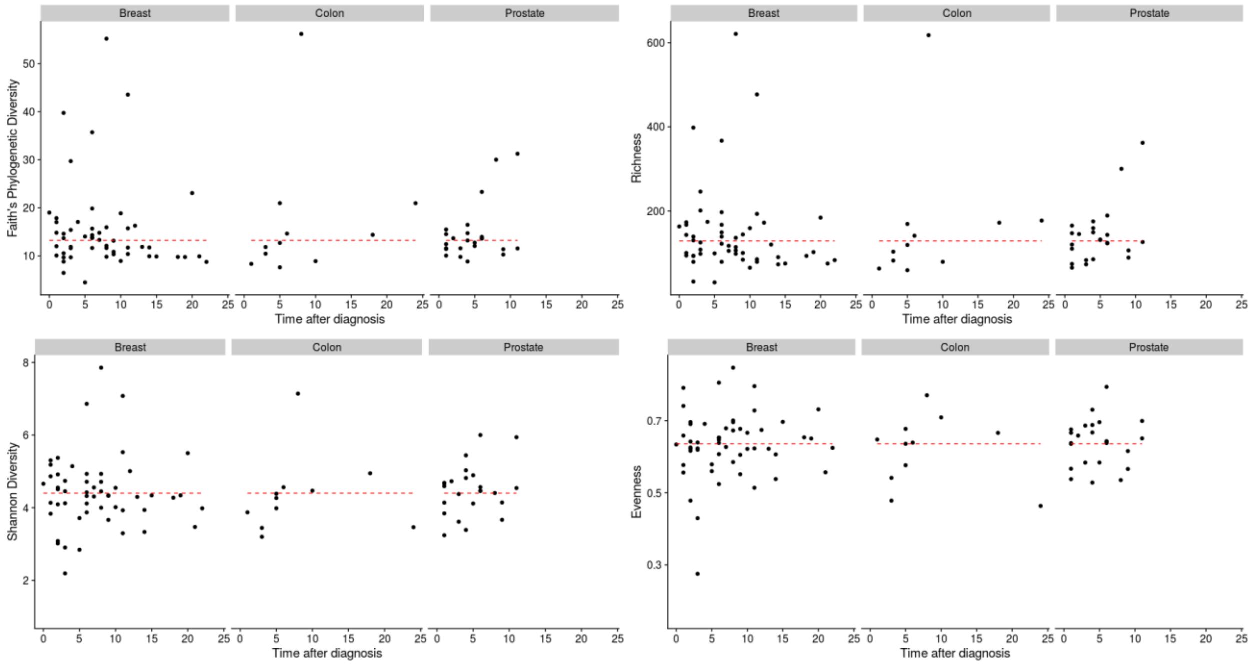


**Supplemental Figure 3. Correlation between alpha diversity and time between cancer diagnosis and sample collection in the retrospective Atlantic PATH cohort.** Within each retrospective cancer type (breast, colon, and prostate cancer) spearman correlation coefficients were calculated between four different alpha diversity metrics (Faith's phylogenetic diversity, richness, Shannon diversity, evenness) and the time between diagnosis and sample collection. We found a significant positive association between richness and time after diagnosis in colon cancer cases (rho=0.62, p=0.04). Red dotted line represents the mean alpha diversity metric among all controls within this cohort.

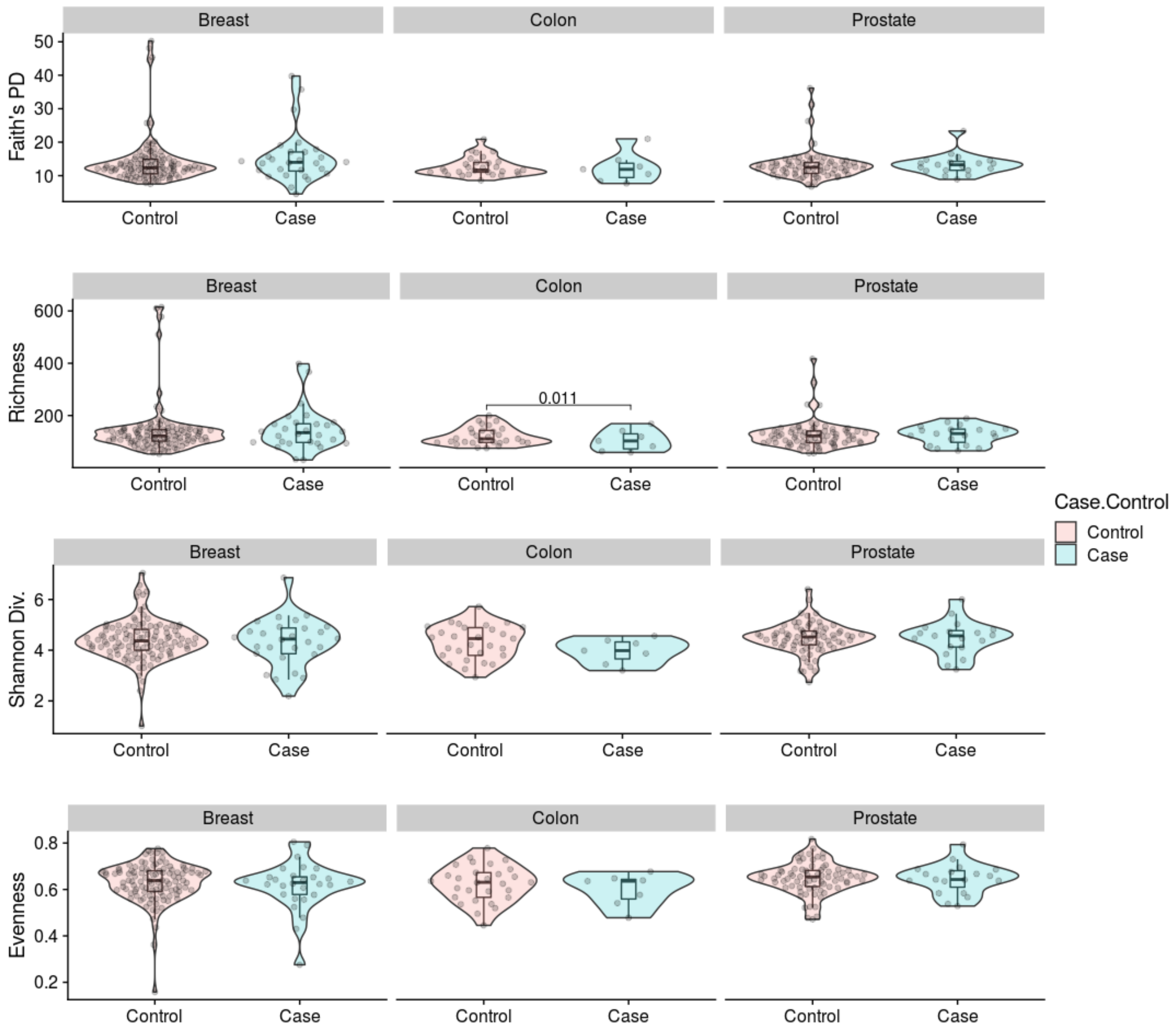


**Supplemental Figure 4**: **Alpha diversity metrics comparing retrospective cases of breast, prostate, and colon cancer within 6 years to matched controls in the retrospective Atlantic PATH cohort.** Case samples were filtered to only include those that were diagnosed within 6 years of sample collection. Each row of boxplots represents a different alpha diversity metric (Faith's phylogenetic diversity, richness, Shannon diversity, evenness) between non-cancer matched controls and retrospective cases of breast, colon, or prostate cancer. We found a significant difference using linear models while controlling for DNA extraction batch in the richness of retrospective colon cancer cases p=0.011. Values above bars represent p-values. Boxplots represent the median and interquartile while violin plots represent the density.

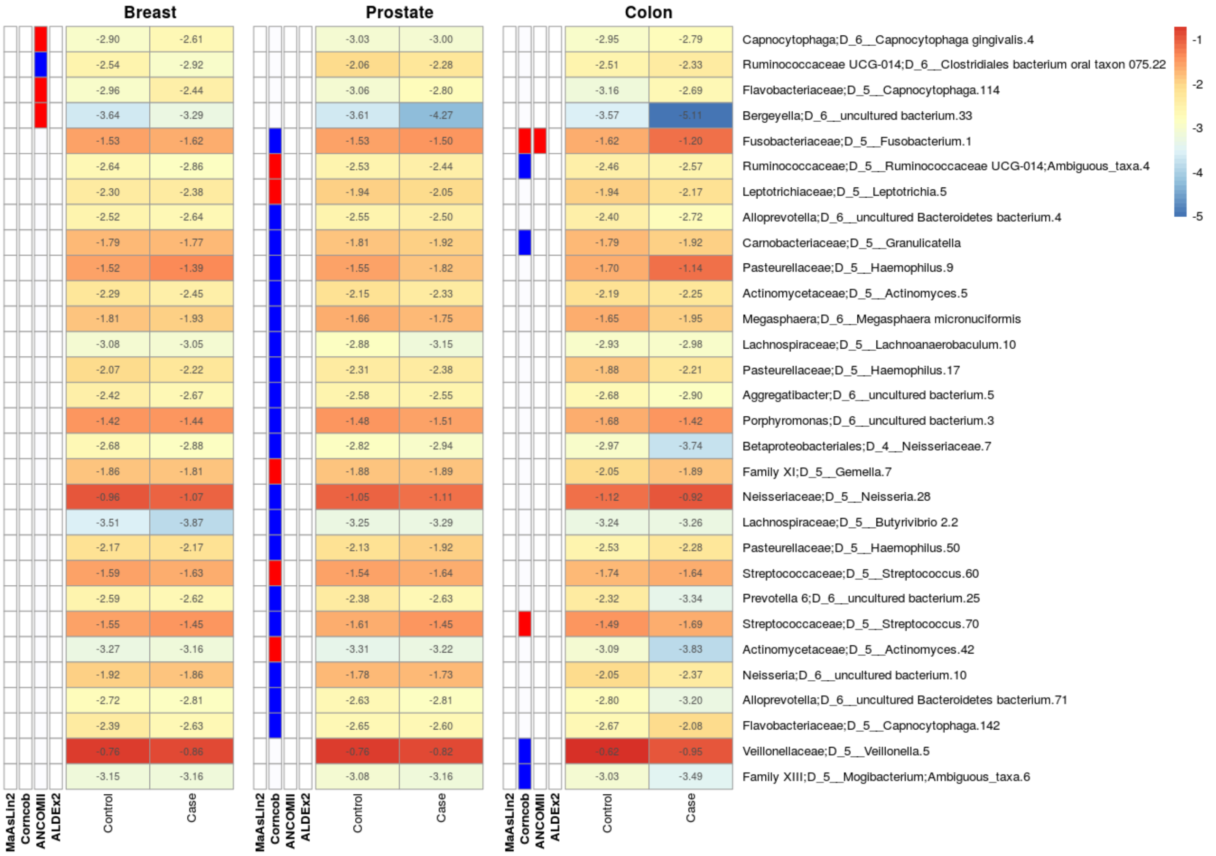


**Supplemental Figure 5. Multiple ASV's are differentially abundant in the oral microbiome of retrospective cases of breast, prostate, and colon cancer in the Atlantic PATH cohort.** The heatmap is divided by cancer type where the first four columns represent the detection of significant associations by one of four tools: MaAsLin2, Corncob, ANCOM-II, and ALDEx2. Blue bars in the first four columns of each subgroup represent a detected increase in control samples while red bars represent a detected increase in case samples. The final two columns within each cancer sub grouping represent the log10 mean relative abundance of each genera with red representing higher abundance values and blue representing lower abundance values.

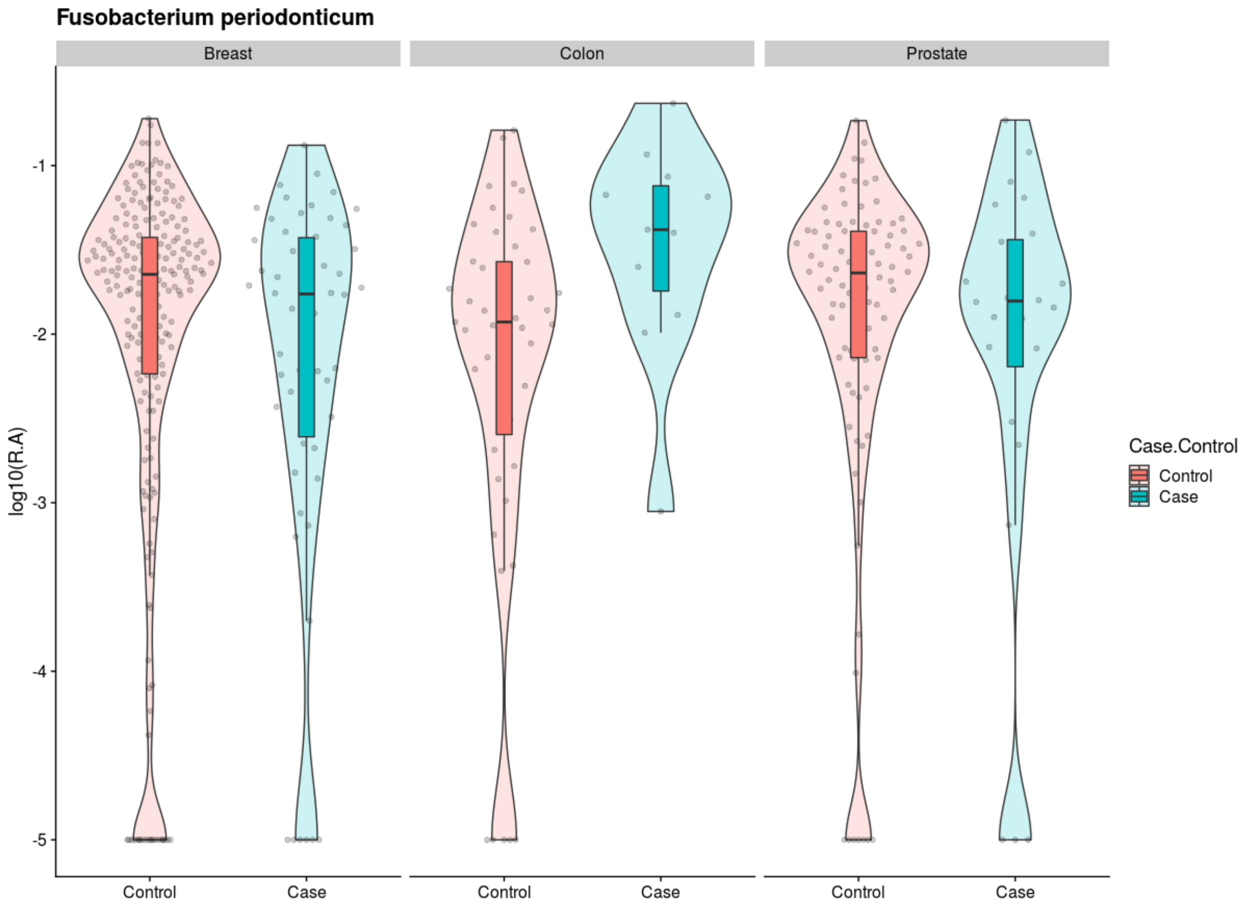


**Supplemental Figure 6. An ASV classified as *Fusobacterium periodonticum* is increased in relative abundance within the oral microbiome of retrospective colon cancer samples in the Atlantic PATH cohort.** Log10 relative abundances of an ASV best classified as *Fusobacterium periodonticum* across retrospective cases of breast, prostate, and colon cancer. Boxplots represent the median and interquartile while violin plots represent the density. A pseudocount of 0.0001 was added before taking log10 of each sample's relative abundance estimate.

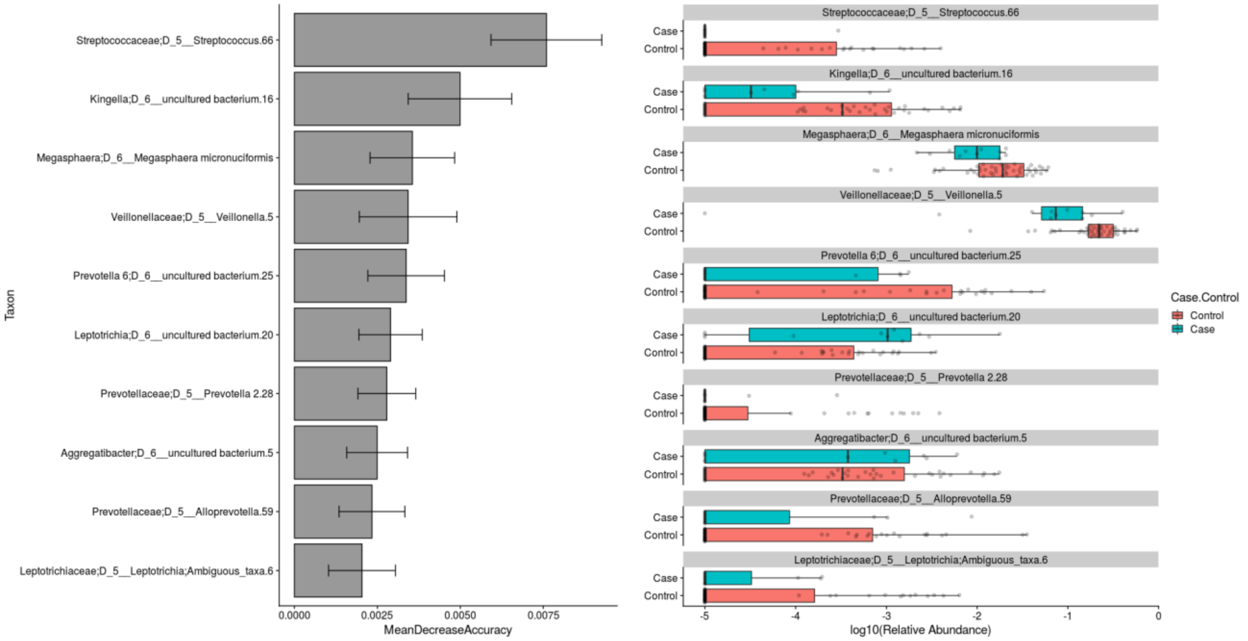


**Supplemental Figure 7. Feature Importance in Random Forest classification of retrospective cases of colon cancer using ASVs normalized to relative abundances in the Atlantic PATH cohort.** MeanDecreaseAccuracy represents feature importance as determined by comparing out-of-bag accuracies before and after permuting the variable of interest across different samples. Features were then sorted by MeanDecreaseAccuracy and the top 10 were plotted along with their log10 relative abundances. A pseudocount of 0.0001 was added before taking log10 of each sample's relative abundance estimate for each ASV. Error bars on bar plots represent standard deviations; boxplots represent the median and inner and outer quartiles, dots within boxplots represent individual datapoints.

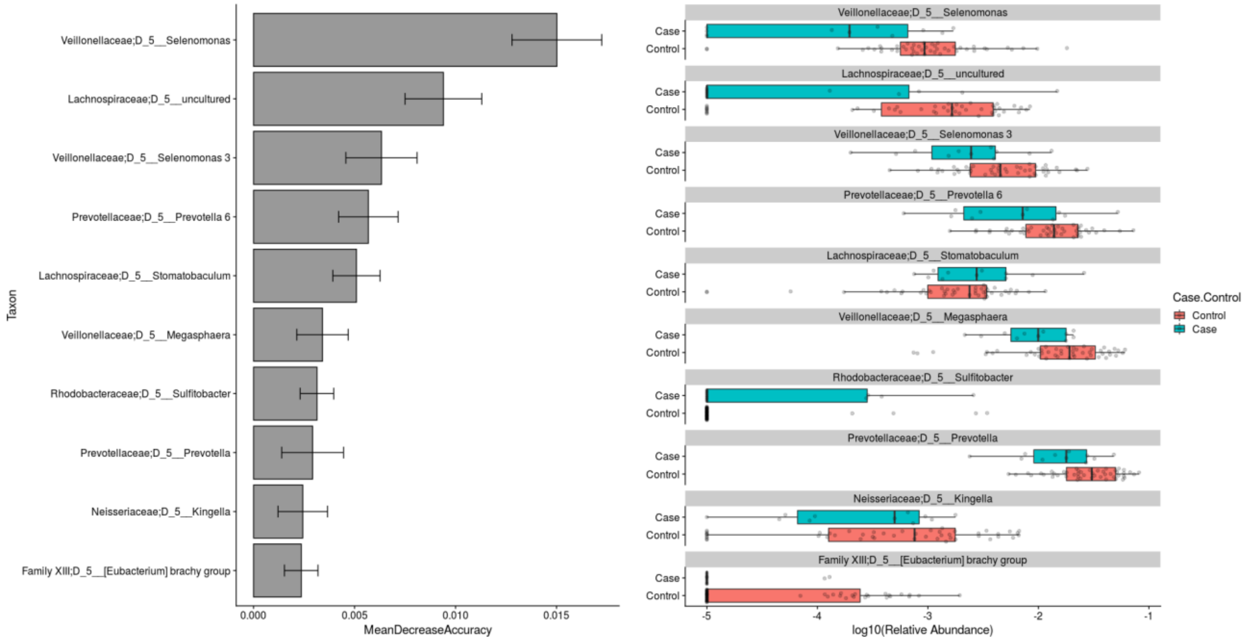


**Supplemental Figure 8. Feature Importance in Random Forest classification of retrospective cases of colon cancer using genera normalized to relative abundances in the Atlantic PATH cohort.** MeanDecreaseAccuracy represents feature importance as determined by comparing out-of-bag accuracies before and after permuting the variable of interest across different samples. Features were then sorted by MeanDecreaseAccuracy and the top 10 were plotted along with their log10 relative abundances. A pseudocount of 0.0001 was added before taking log10 of each sample's relative abundance estimate for each genus. Error bars on bar plots represent standard deviations; boxplots represent the median and interquartile; dots within boxplots represent individual datapoints.

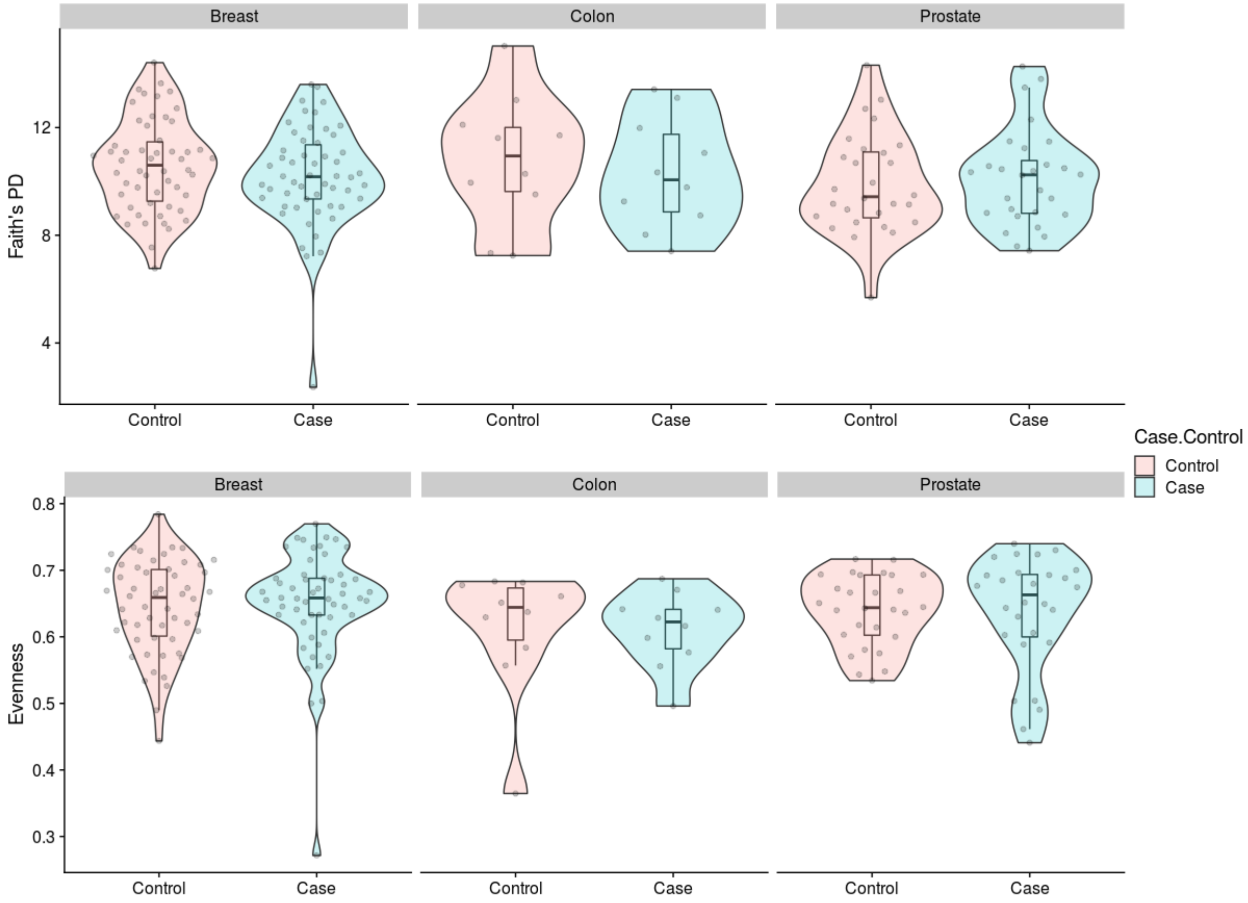


**Supplemental Figure 9. Comparison of alpha diversity between prospective cancer cases and non-cancer match controls in the Atlantic PATH cohort.** Faith's phylogenetic diversity and evenness were compared within the PATH cohort between non-cancer matched controls and prospective cases of breast, colon, and prostate cancer. Each alpha diversity metric was compared within each cancer type using linear models. No significant differences were detected. Boxplots represent the median and interquartile while violin plots represent the density.

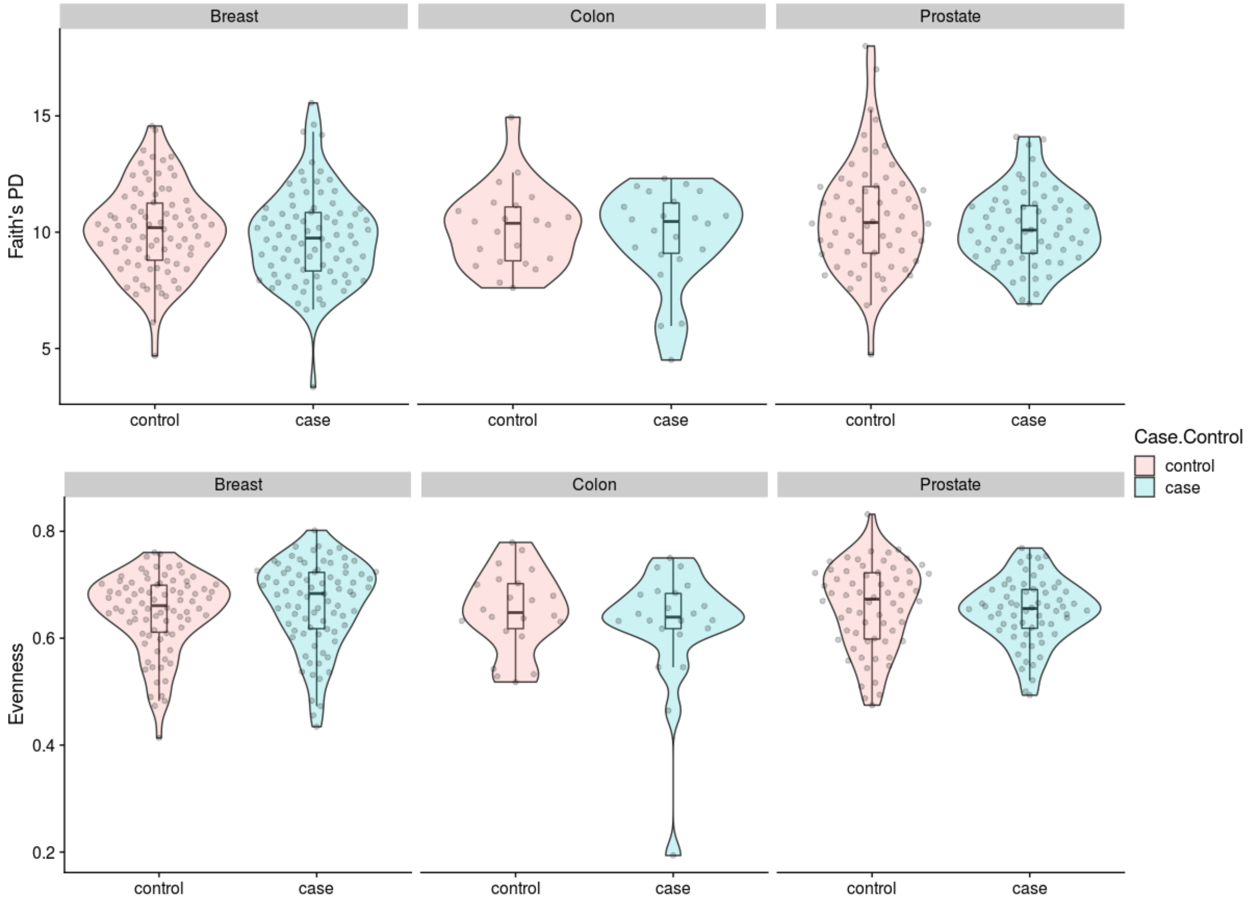


**Supplemental Figure 10. Comparison of alpha diversity between prospective cancer cases and non-cancer match controls in the ATP cohort** . Faith's phylogenetic diversity and evenness were compared within the ATP cohort between non-cancer matched controls and prospective cases of breast, colon, and prostate cancer. Each alpha diversity metric was compared within each cancer type using linear models. No significant differences were detected. Boxplots represent the median and interquartile while violin plots represent the density.

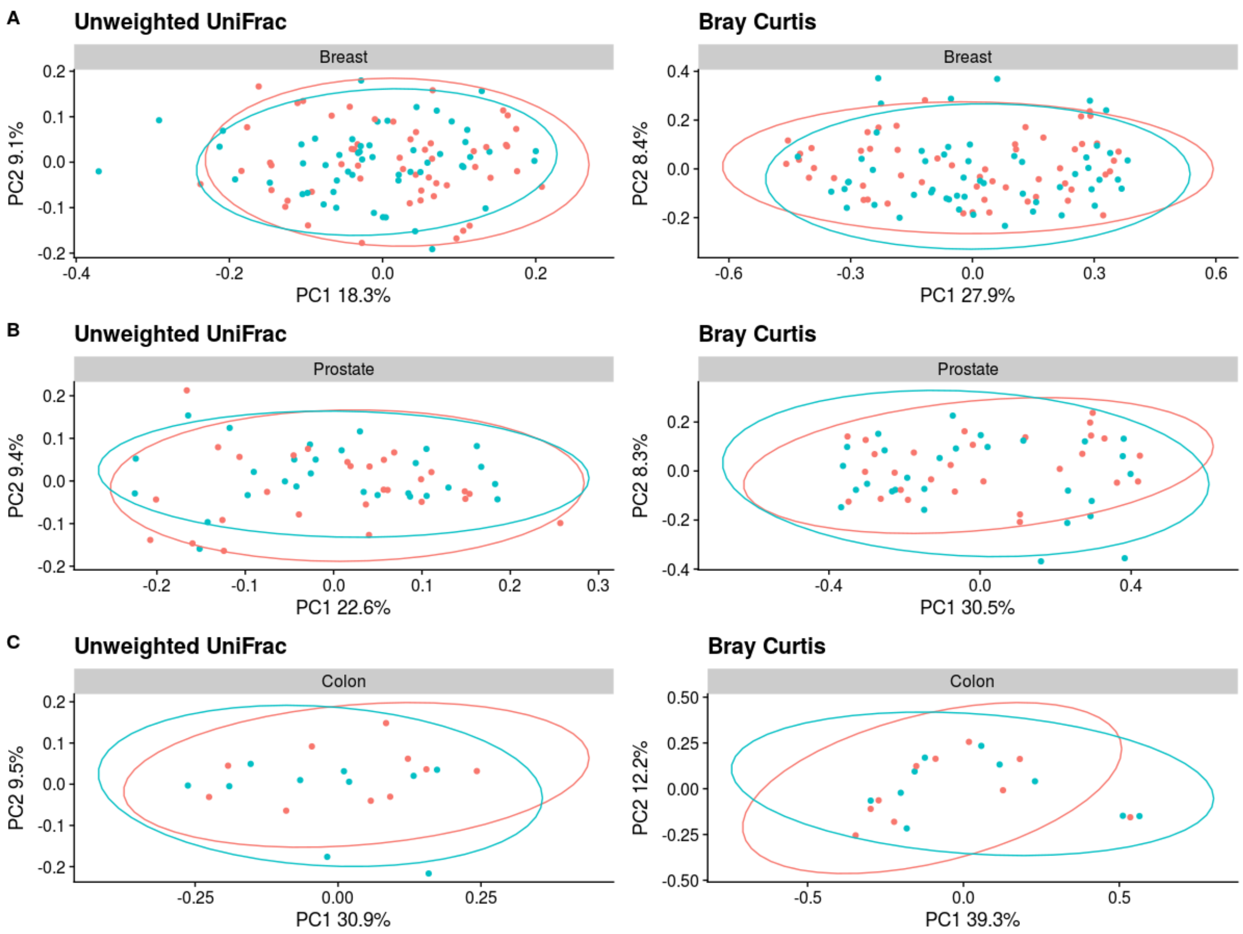


**Supplemental Figure 11. Case-Control beta diversity analysis of prospective cases of breast, prostate, or colon cancer in the Atlantic PATH cohort.** Two different beta diversity metrics (unweighted UniFrac and Bray-Curtis dissimilarity) within the PATH cohort were compared between non-cancer matched controls and breast, prostate or colon cancer. No significant differences were found within each metric and cancer type using a PERMANOVA test.

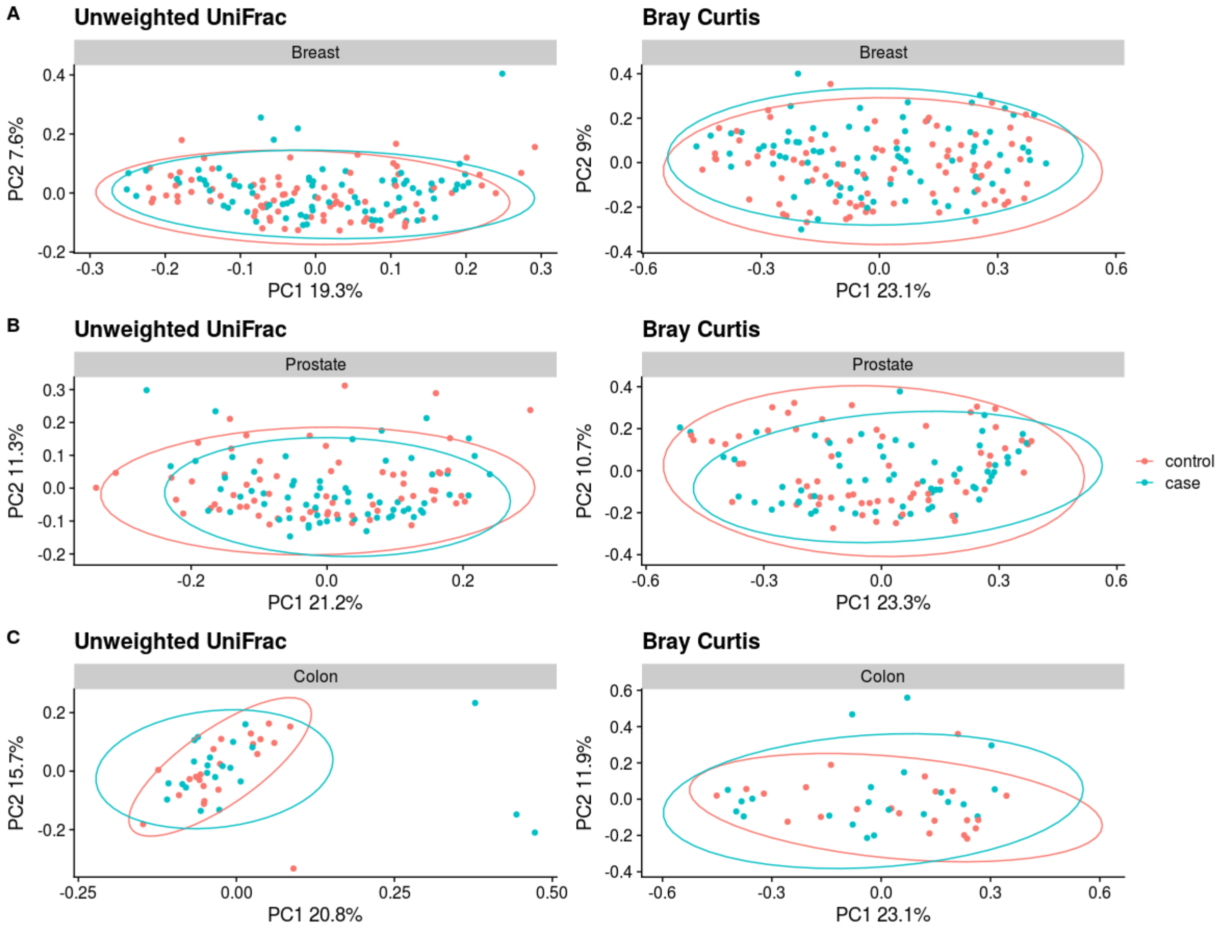


**Supplemental Figure 12. Case-Control beta diversity analysis of prospective cases of breast, prostate, or colon cancer in the ATP cohort.** Two different beta diversity metrics (unweighted UniFrac and Bray-Curtis dissimilarity) within the ATP cohort were compared between non-cancer matched controls and breast, prostate or colon cancer. No significant differences were found within each metric and cancer type using a PERMANOVA test.

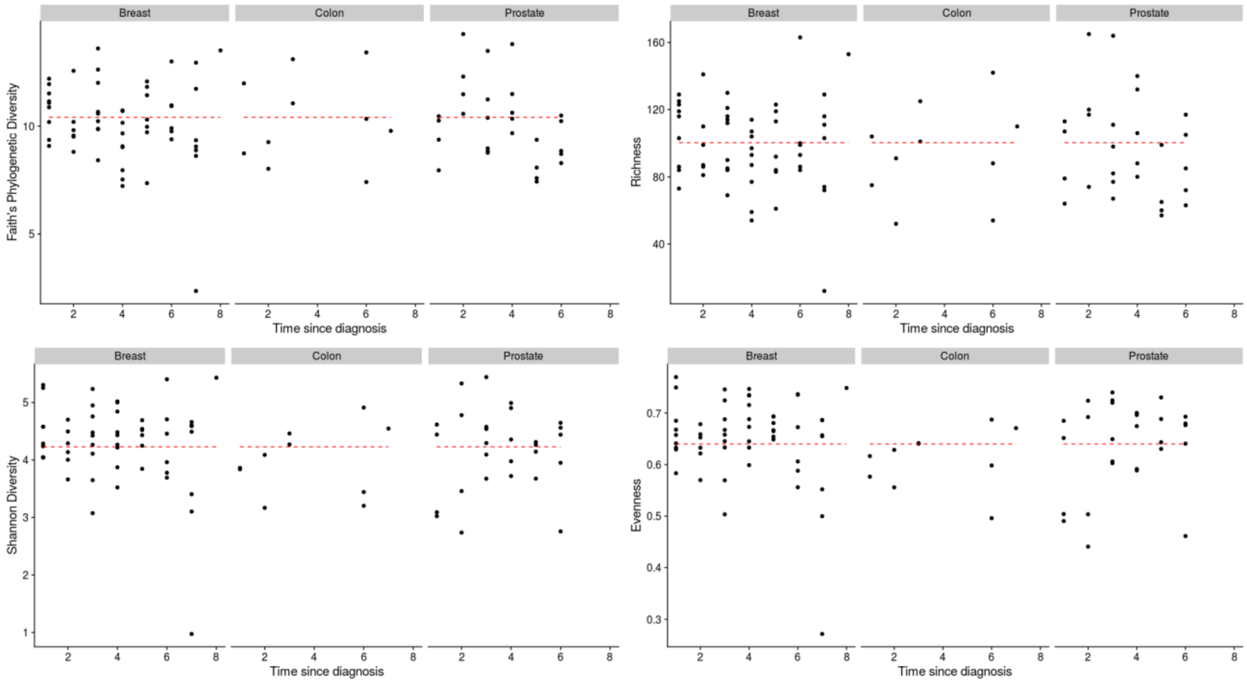


**Supplemental figure 13. Correlation between alpha diversity and time since diagnosis in prospective cancer cases within the Atlantic PATH cohort.** Within the cohort prospective cancer cases were divided into type (breast, colon, and prostate cancer) and spearman correlations were calculated between four different alpha diversity metrics (Faith's phylogenetic diversity, richness, Shannon diversity, evenness) and the time between sample collection and disease diagnosis. No significant correlations were found.

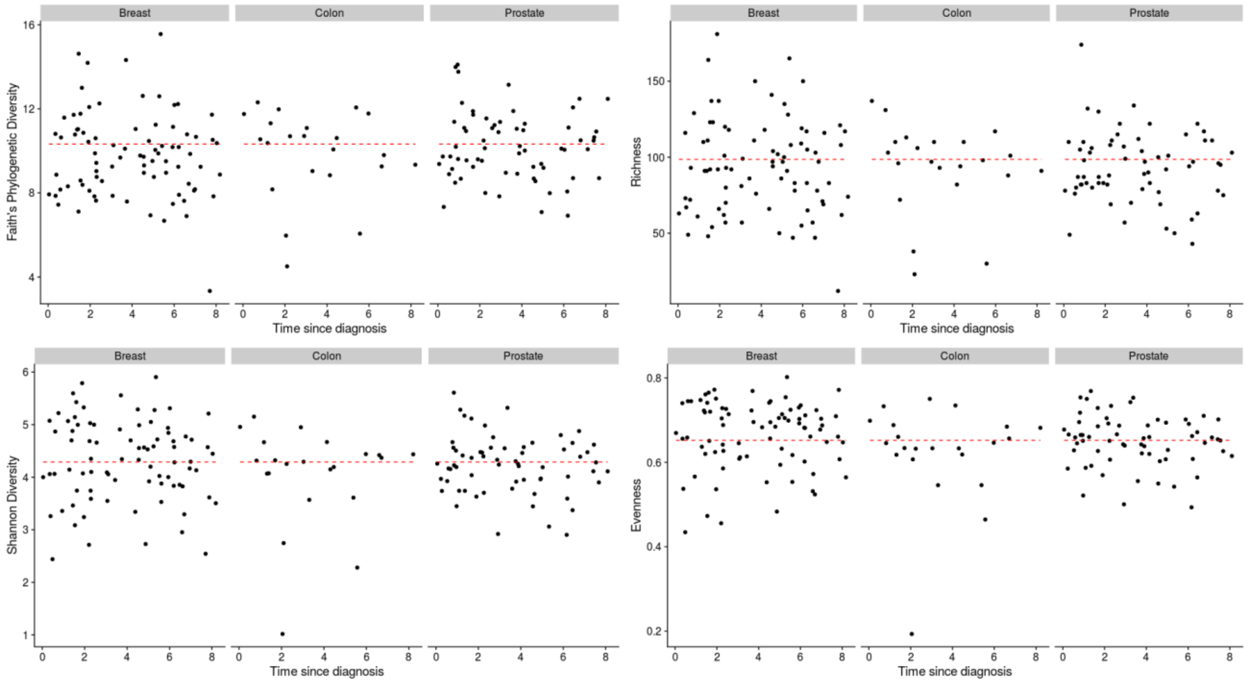


**Supplemental Figure 14. Correlation between alpha diversity and time since diagnosis in prospective cancer cases within the ATP cohort.** Within this cohort prospective cancer cases were divided into type (breast, colon, and prostate cancer) and spearman correlations were calculated between four different alpha diversity metrics (Faith's phylogenetic diversity, richness, Shannon diversity, evenness) and the time between sample collection and disease diagnosis. No significant correlations were found.

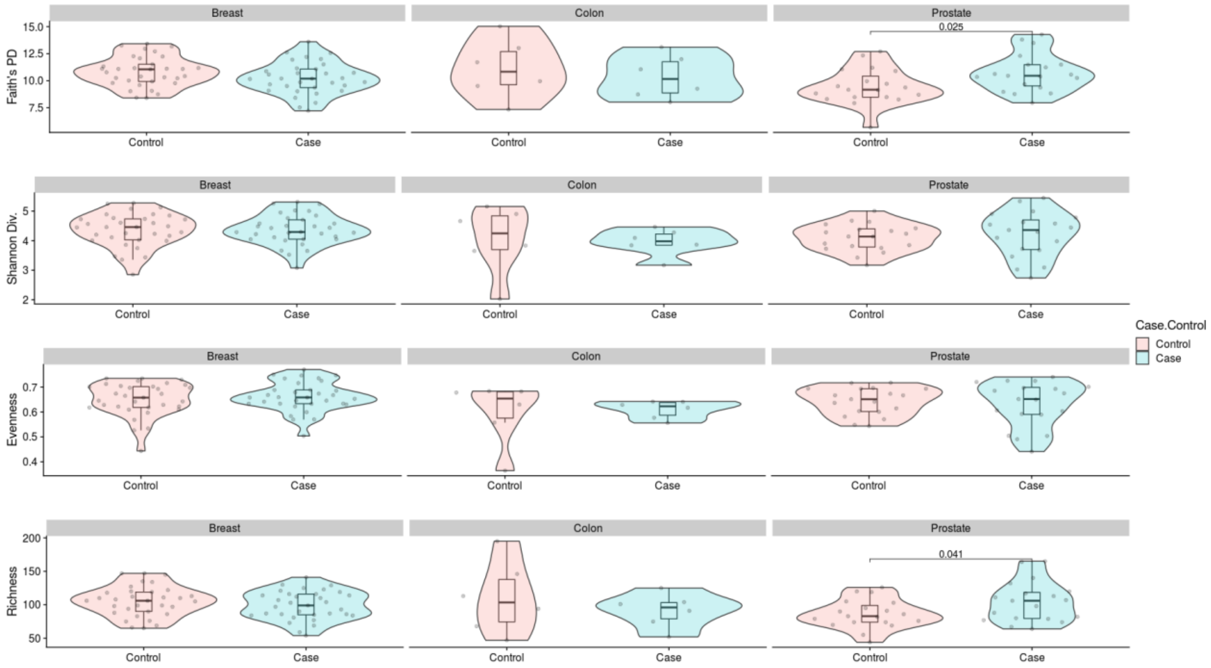


**Supplemental Figure 15. Alpha diversity metrics comparing prospective cases of breast, prostate, and colon cancer within 4 years to matched non-cancer controls in the Atlantic PATH cohort.** Prospective case samples within the cohort samples were filtered to only include those that were diagnosed within 4 years of sample collection. Four different alpha diversity metrics (Faith's phylogenetic diversity, richness, Shannon diversity, evenness) were compared between non-cancer matched controls and prospective cases of breast, colon, or prostate cancer. We found a significant difference using linear models in the richness and Faith's phylogenetic diversity in prospective prostate cancer samples. Values above bars present p-values. Boxplots represent the median and interquartile while violin plots represent the density.

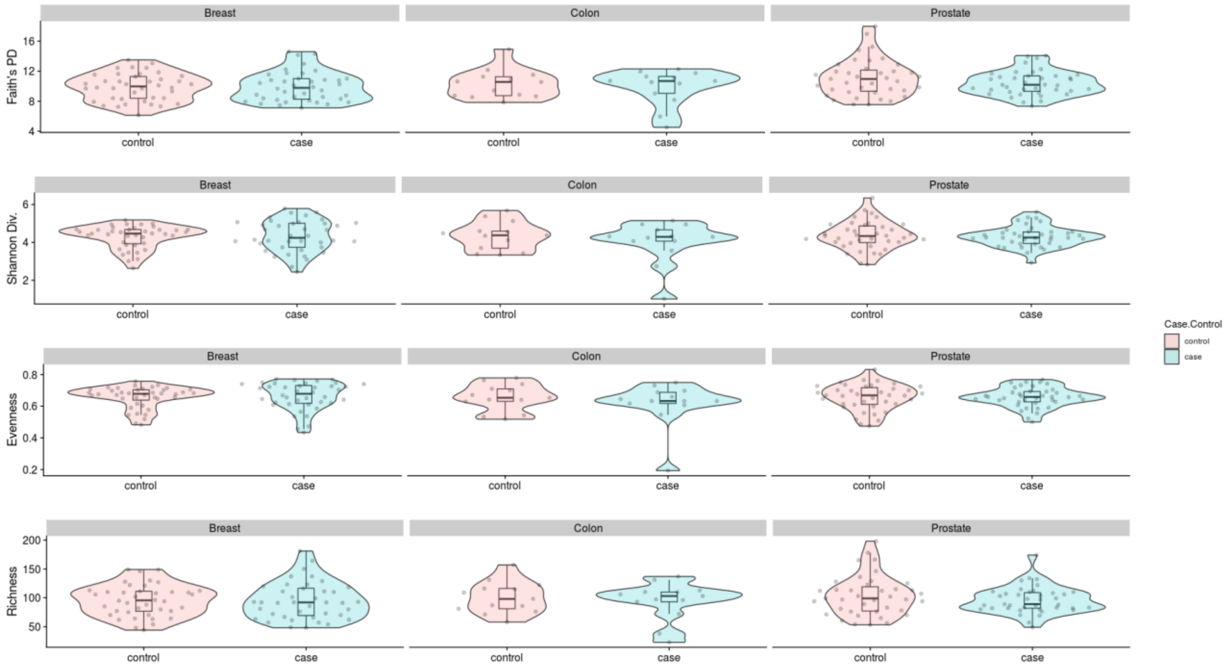


**Supplemental Figure 16. Alpha diversity metrics comparing prospective cases of breast, prostate, and colon cancer within 4 years to matched non-cancer controls in the ATP cohort.** Prospective case samples with the ATP cohort were filtered to only include those that were diagnosed within 4 years of sample collection. Four different alpha diversity metrics (Faith's phylogenetic diversity, richness, Shannon diversity, evenness) were compared between non-cancer matched controls and prospective cases of breast, colon, or prostate cancer. No significant differences were found using linear models comparing case and control samples. Boxplots represent the median and interquartile while violin plots represent the density.

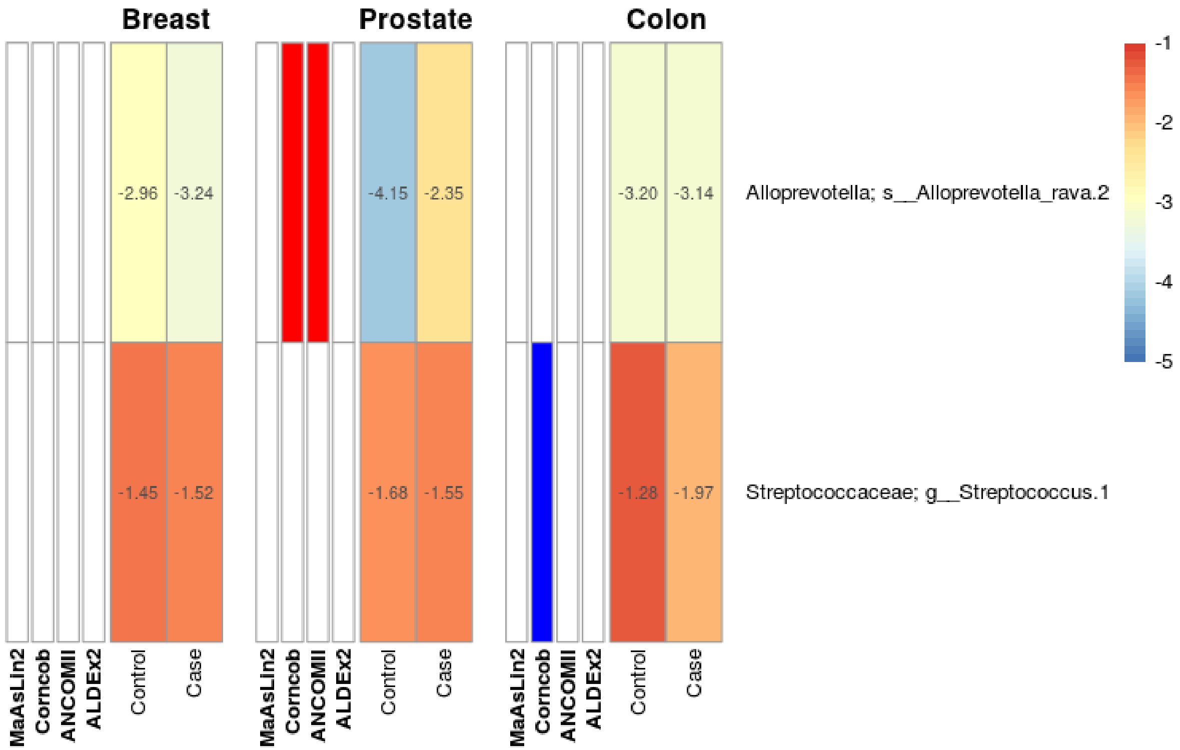


**Supplemental Figure 17. Two ASVs were detected as being differentially abundant in the oral microbiome of prospective cases of breast, prostate, and colon cancer in the Atlantic PATH cohort.** The heatmap is divided by cancer type where the first four columns represent the detection of significant associations by one of four tools: MaAsLin2, Corncob, ANCOM-II, and ALDEx2. Blue bars in the first four columns of each subgroup represent a detected increase in control samples while red bars represent a detected increase in case samples. The final two columns within each cancer sub grouping represent the log10 mean relative abundance of each ASV with red representing higher relative abundance values and blue representing lower relative abundance values. One ASV, classified as *Alloprevotella rava*, was detected by both corncob and ANCOM-II in prospective prostate cancer cases. Additionally, one ASV, classified at the genus level as Streptococcus, was also detected by corncob in prospective colon cancer cases.

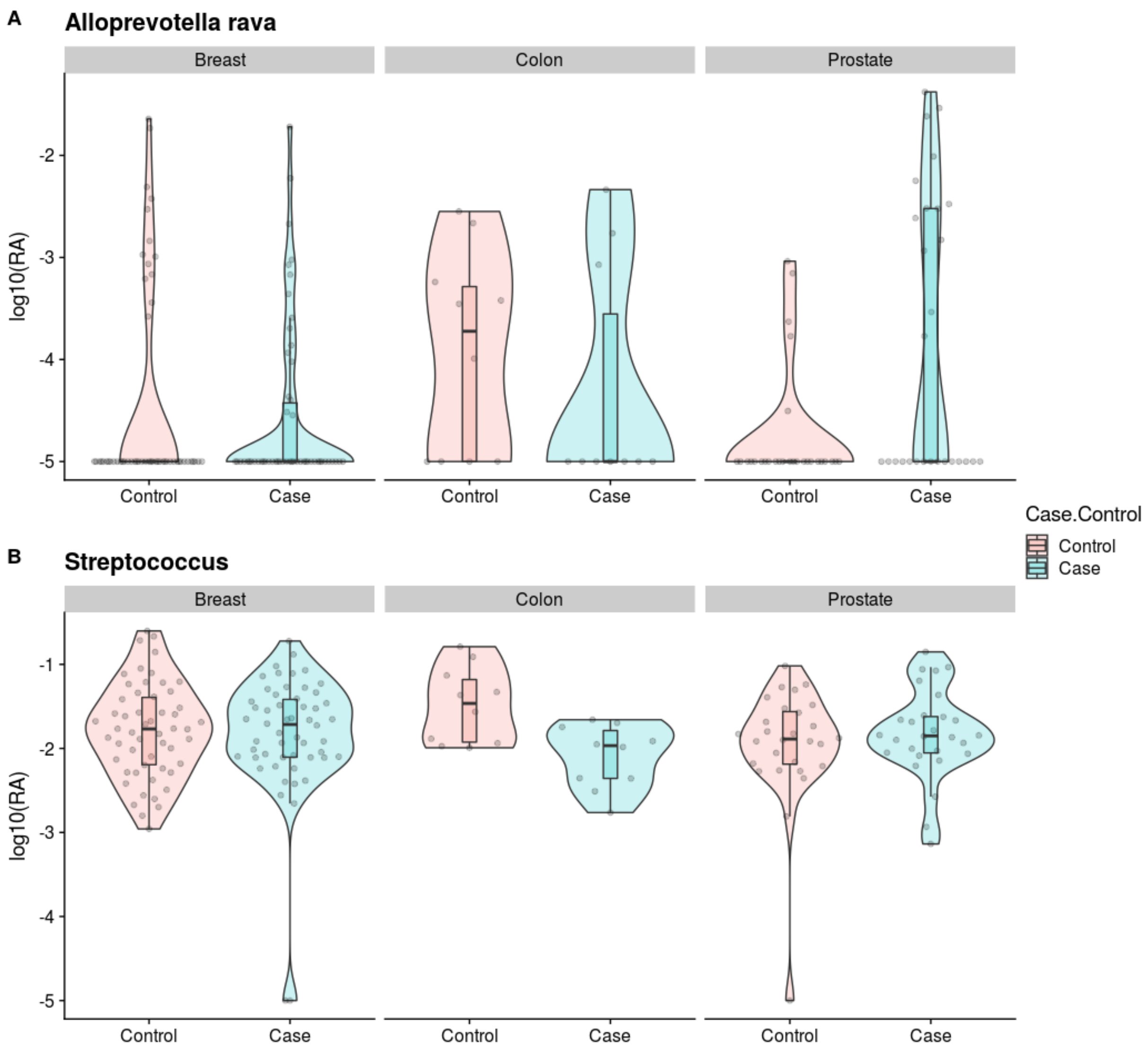


**Supplemental Figure 18. ASVs detected as differentially abundant in prospective cases of prostate and colon cancer within the Atlantic PATH cohort.** Log10 relative abundance values with a pseudo count of 0.0001 compared between case and control samples for the two ASVs detected as being differentially abundant in prospective cases of prostate cancer (A) and colon cancer (B). Boxplots represent the median and interquartile while violin plots represent the density.

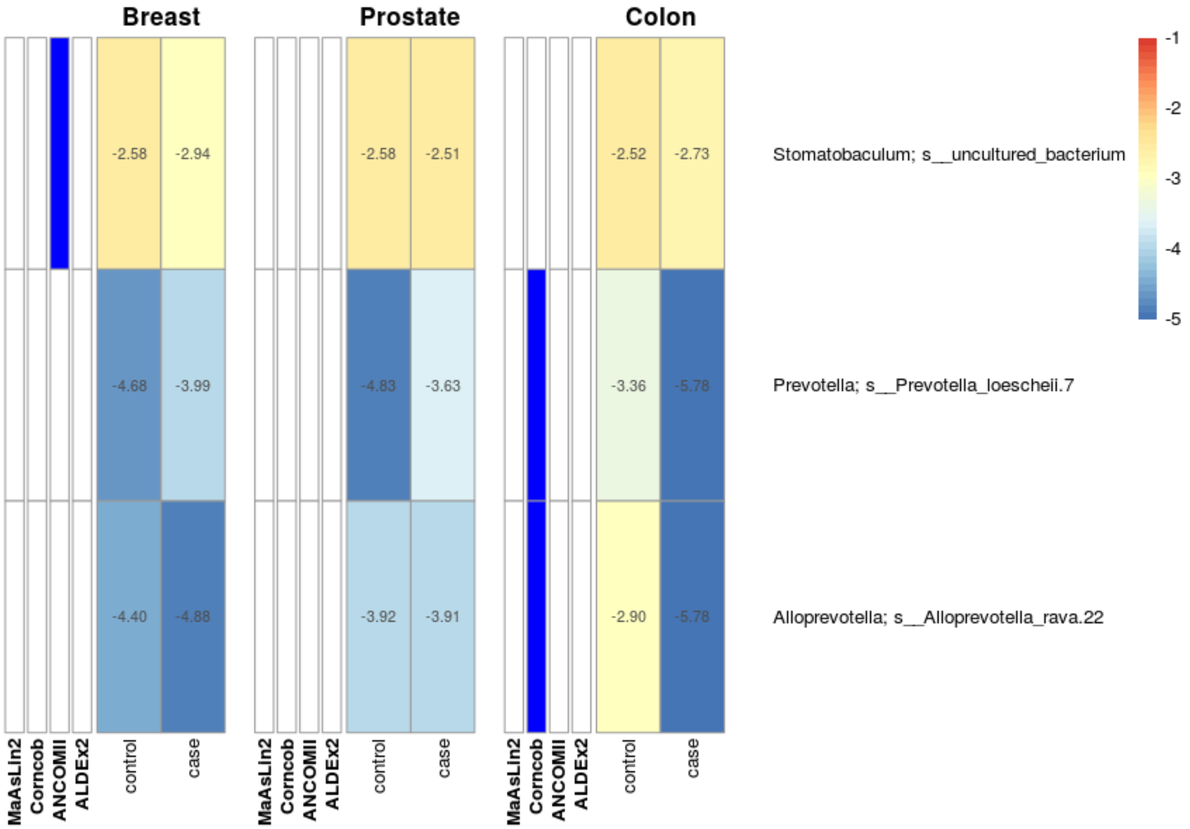


**Supplemental Figure 19. Two ASVs were detected as being differentially abundant in the oral microbiome of prospective cases of colon cancer in the ATP cohort.** The heatmap is divided by cancer type where the first four columns represent the detection of significant associations by one of four tools: MaAsLin2, Corncob, ANCOM-II, and ALDEx2. Blue bars in the first four columns of each subgroup represent a detected increase in control samples while red bars represent a detected increase in case samples. The final two columns within each cancer sub grouping represent the log10 mean relative abundance of each ASV with red representing higher relative abundance values and blue representing lower relative abundance values. Two ASVs, classified as *Prevotella loescheii* and *Alloprevotella rava* were detected by corncob to be decreased in relative abundance in colon cancer. One ASV classified as an uncultured species of Stomatobaculum was detected by corncob to be decreased in relative abundance in breast cancer.

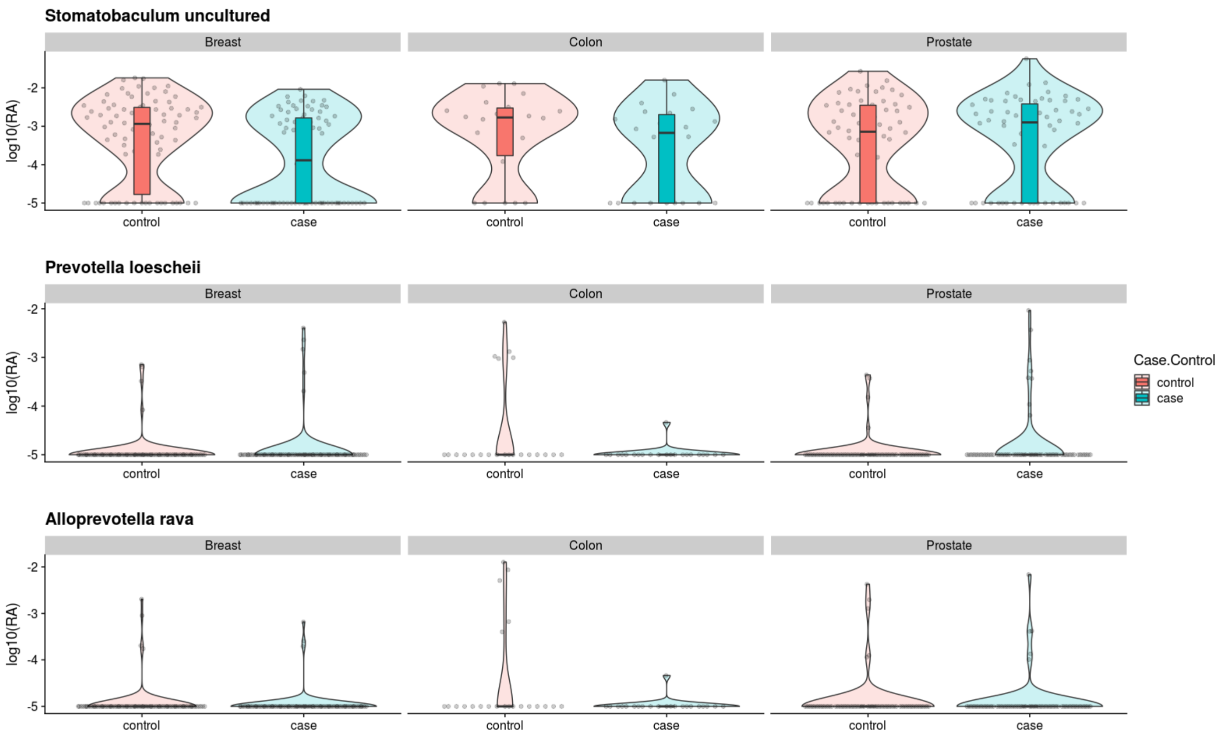


**Supplemental Figure 20. ASVs detected as differentially abundant in prospective cases of colon cancer and breast cancer within the ATP cohort.** Log10 relative abundance values with a pseudo count of 0.0001 compared between case and control samples for the two ASVs detected as being differentially abundant by corncob in prospective cases of colon cancer. One ASV detected as being differentially abundant by corncob in prospective cases of breast cancer. Boxplots represent the median and interquartile while violin plots represent the density.

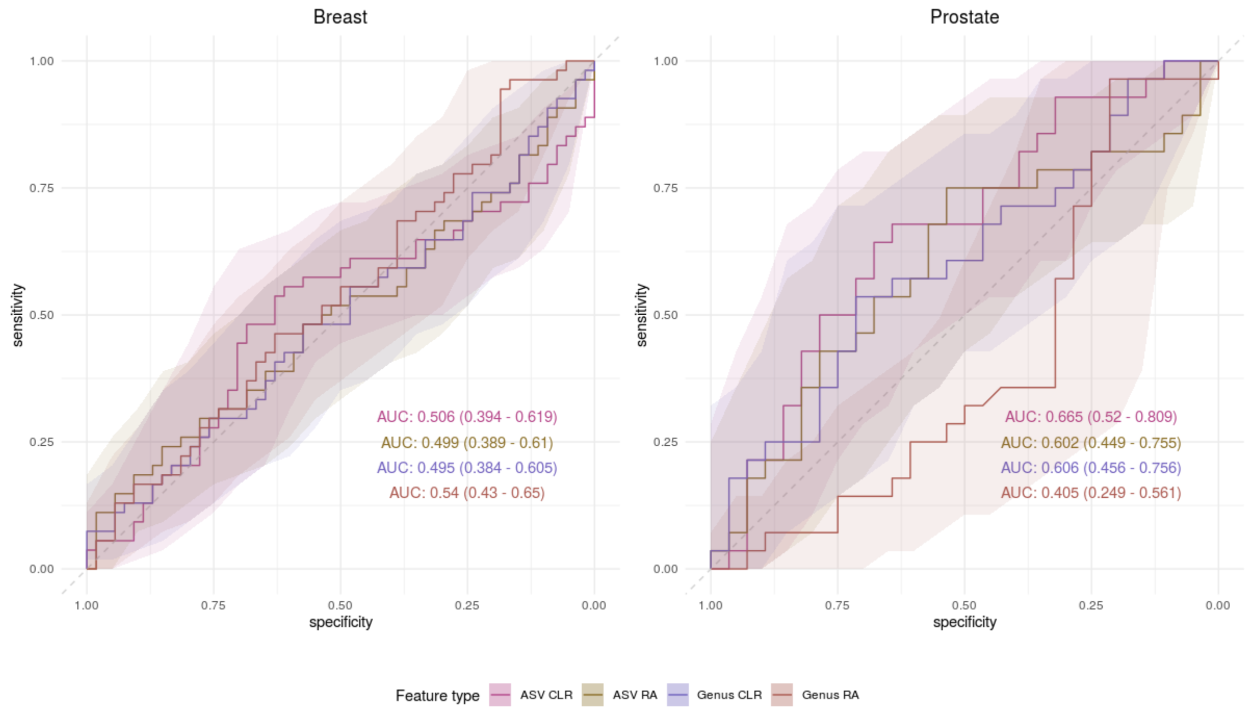


**Supplemental Figure 21. Random Forest classification of prospective cases of breast, prostate, and colon cancer based on microbial taxonomic composition within the Atlantic PATH cohort.** Receiver operator curves (ROC) showing the specificity and sensitivity of the classification of non-cancer matched controls and prospective cases of breast, prostate, or colon cancer within the PATH cohort. Models were constructed using 100-repeat 5-fold cross validation and hold-out performance was determined through taking the mean number of votes for each hold-out sample across all 100 repeats. Within each plot four different ROCs are represented showing the classification accuracy using ASVs or genera normalized with either total-sum-scaling or center-log-ratio abundance. Shaded areas represent 95% confidence intervals determined through 2000 bootstrap samplings.

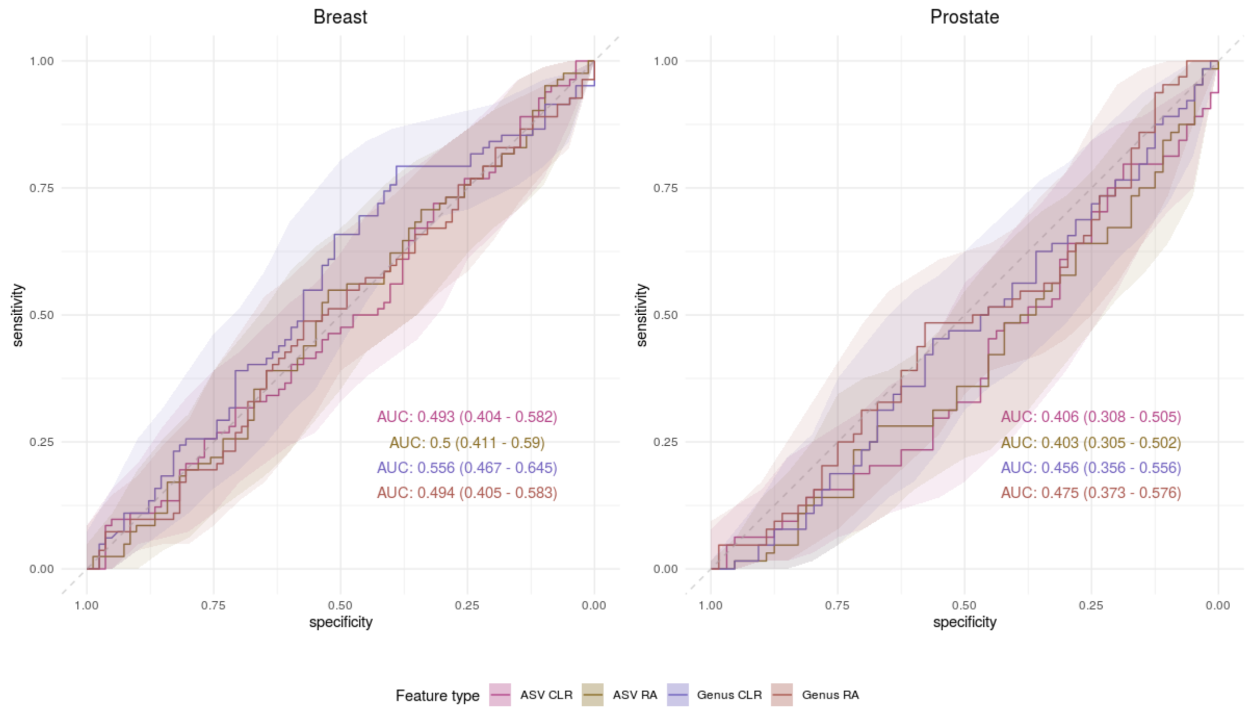


**Supplemental Figure 22. Random Forest classification of prospective cases of breast, prostate, and colon cancer based on microbial taxonomic composition within the ATP cohort.** Receiver operator curves (ROC) showing the specificity and sensitivity of the classification of non-cancer matched controls and prospective cases of breast, prostate, or colon cancer within the ATP cohort. Models were constructed using 100-repeat 5-fold cross validation and hold-out performance was determined through taking the mean number of votes for each hold-out sample across all 100 repeats. Within each plot four different ROCs are represented showing the classification accuracy using ASVs or genera normalized with either total-sum-scaling or center-log-ratio abundance. Shaded areas represent 95% confidence intervals determined through 2000 bootstrap samplings.
